# Integration of sexual dimorphism and left-right asymmetry in the development of the duck syrinx

**DOI:** 10.1101/2024.12.10.627783

**Authors:** Darcy Mishkind, ChangHee Lee, Chaitra Prabhakara, Penelope Tir, Evan P. Kingsley, Julia A. Clarke, Franz Goller, Clifford J. Tabin

## Abstract

Embryonic morphogenesis is regulated across multiple dimensions. In ducks, the syrinx, the avian vocal organ, undergoes morphogenic processes that result in both left-right asymmetric and sexually dimorphic development. Although these properties are thought to be controlled by the NODAL-PITX2 left-right signaling cascade and the sex steroid pathways, how these mechanisms work together to produce an asymmetric structure in a sexually dimorphic manner remains unclear. Here, we first establish evidence for sexual selection driving the evolution of the duck syrinx. During its development, we observe that *PITX2* is expressed on the left side in both male and female ducks, although not in other birds. Asymmetric activation of *PITX2* in this domain is triggered by bilateral BMP signaling but is limited to the left side by left-specific stably accessible chromatin established during an earlier asymmetric wave of *PITX2* expression. Ultimately, there is an induction of left-specific WNT and BMP signaling in the syrinx primordium, which synergistically elevates cell proliferation on the left, leading to asymmetric growth. Estrogen receptor expression is also shown to be induced on the left side of the forming syrinx. This has no effect in males, but in females where the hormone is present, estrogen signaling reduces left-sided cell proliferation, thus promoting bilaterally symmetric growth. These data demonstrate how sexually dimorphic left-right asymmetry can be integrated to produce an adaptive trait.

## Introduction

The syrinx is the unique vocal organ of birds, located at the tracheal-bronchial junction. Though the syrinx shows striking morphological diversity, allowing it to produce the wide array of sounds generated by different bird species, in most species it is bilaterally symmetric^1^. For example, the chick (*Gallus gallus*), a member of the clade Galliformes (chickens, turkey, grouse, etc.), shows no asymmetry in the syrinx^2^. However, in the closely related family Anatidae (ducks, geese, and swans) many species show a syringeal asymmetry that is moreover sex-specific; females develop a bilaterally symmetric syrinx, while males develop a laterally asymmetric syrinx with a left-sided hollow, initially cartilaginous outgrowth, or bulla^3–7^. Lateral asymmetry in the syrinx of ducks and their relatives may date back to the Mesozoic Era and be specific to this clade within the broader Anseriformes; the fossil of a relative of Anatidae, *Vegavis iaai*, was found to have asymmetric bronchi, while the basal Anseriform *Presbyornis* showed no asymmetry^8^. The bulla has been proposed to be a resonance chamber that allows male ducks to produce whistling vocalizations during courtship, although a direct link between bulla morphology and the frequencies produced during vocalization has not been demonstrated^7,9^.

From a developmental perspective, the formation of a left-right asymmetric, sexually dimorphic structure presumably requires the input of two distinct signaling pathways, the NODAL cascade known to be responsible for left-right asymmetric morphogenesis and the sex steroid pathways responsible for sex-specific morphogenesis. Yet how these signals direct the excessive growth of the cartilage on the left side of the male syrinx to form a bulla remains unclear, as does the mechanism by which NODAL signaling and sex steroid activity are ultimately integrated during syrinx morphogenesis.

While left-right asymmetric morphogenesis has never been addressed at a molecular level in the context of the syrinx, a known set of factors play a highly conserved role in the determination of morphological left-right asymmetry throughout metazoa^10–13^. The left-right asymmetry cascade is initiated during gastrulation with the activation of the transforming growth factor-β (TGFβ)-related signal, NODAL, throughout the left side of the lateral plate mesoderm (LPM). NODAL acts upstream of the transcription factor PITX2^14–16^. Genetic studies have indicated that PITX2, in turn, is responsible for directing subsequent left-right asymmetric development of the internal organs^13,17,18^. Consistent with this, *PITX2* expression can be observed on the left side of tissues undergoing asymmetric morphogenesis. For example, this gene is specifically expressed in the left side of the dorsal mesentery during asymmetric gut looping^19,20^. Interestingly, recent work has progressed our understanding of *PITX2* expression dynamics; during the development of the chick gut, the initial left-sided *PITX2* expression in the LPM subsides before asymmetric *PITX2* expression reemerges in a second wave within this LPM-derived tissue, and ultimately induces asymmetric gut rotation^21^. However, the activity of *PITX2* during asymmetric development of the duck syrinx has not been explored.

In addition to being left-right asymmetric, the duck syrinx is sexually dimorphic. In principle, a male-specific structure could be induced only in males through the action of androgens in male embryos, or alternatively by being repressed specifically in females through the action of estrogens. Previous studies have shown that the latter is the basis for the dimorphism of the duck syrinx; these works observed that the duck syrinx develops a male-type bulla when grown in the absence of hormones, regardless of sex^5,6^. In addition, estrogen treatment yields symmetric bulla-less male syrinxes^3^. In the current study, we explore how the estrogen pathway intersects with the NODAL-PITX2 cascade to produce a left-right asymmetric bulla exclusively in the syrinx of male ducks.

## Results

A striking feature of the syrinx in the adult duck is the large osseous bulla on the left side present only in males (Figure 1a,b). This structure has been hypothesized to function in producing male-specific mating calls, although no direct evidence for this has yet been described. In particular, the bulla has been suggested to make whistling sounds as a Helmholtz resonator, in which air forced into a cavity vibrates at a specific frequency determined by the geometry of the cavity (as happens when one blows air across the top of a bottle)^7^. To determine the inner geometry of a mallard duck bulla, an adult syrinx was imaged by micro-computed tomography (µCT) and inner dimensions were measured (Movie S1; Table 1). The fundamental frequency was calculated using the Helmholtz equation. This predicted frequency was then compared to frequencies analyzed from recordings of vocalizing male mallards, publicly available from Cornell’s Macaulay Library ornithology database. Strikingly, the predicted fundamental frequency produced by the bulla matched the peak frequency (in this case also the fundamental frequency) of the whistle made during male-specific courtship vocalizations (Figure 1c). In contrast, the peak frequencies emitted by male mallards in non-mating vocalization did not match the predicted frequency of the bulla (Figure 1c). A recording capturing a high frequency vocalization as well as pressure data shows the whistle being produced on the inspiration (when pressure in the bulla drops) rather than the exhalation, which may be how the courtship vocalization is produced (Figure S1). These data support a functional role for the bulla in Anatidae courtship behavior. This in turn suggests that the evolution of the asymmetric, sexually dimorphic duck syrinx may have been driven by sexual selection.

**Figure 1.**
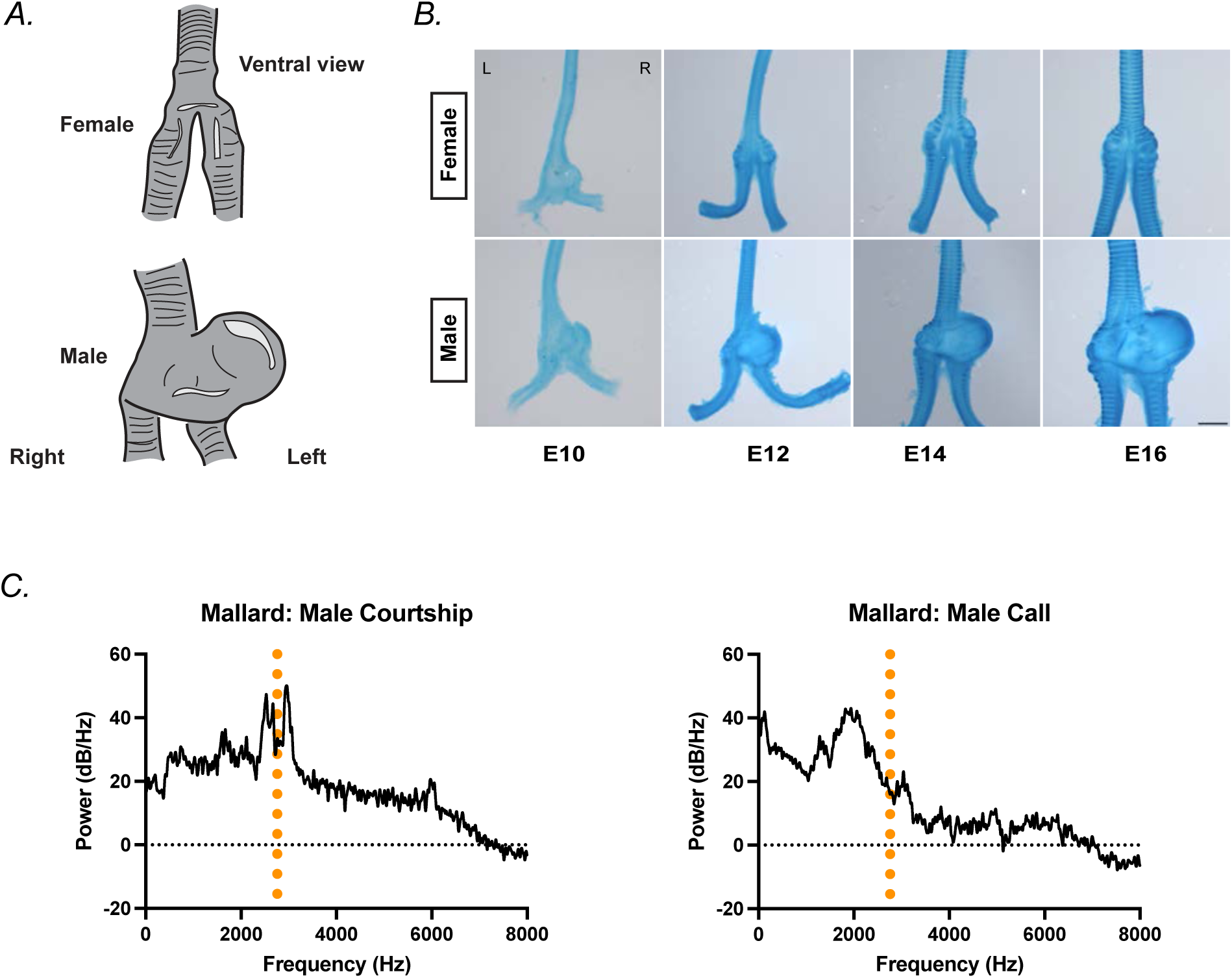
Overview of the duck syrinx form and evidence that the bulla may allow males to produce courtship vocalizations. (A) The syrinxes of the male and female adult mallard and khaki campbell show clear sexual dimorphism. Female after Yilmaz et al., 2012, male after Frank et al., 2007 and dissected Khaki Campbell specimen^1,2^. (B) Alcian blue stained syrinxes from embryonic time points that span the divergence of female (above) and male (below) syringeal development from embryonic day 10 (E10) to embryonic day 16 (E16). Ventral facing up. Right (R) and left (L) as labeled on image. Scale bar = 1mm. (C) Frequencies predicted to be produced by the male bulla modeling it as a Helmholtz resonator (orange dotted line) as compared to power spectra of wild recordings in the mallard, left, male courtship call, right, male call.

**Table 1.**
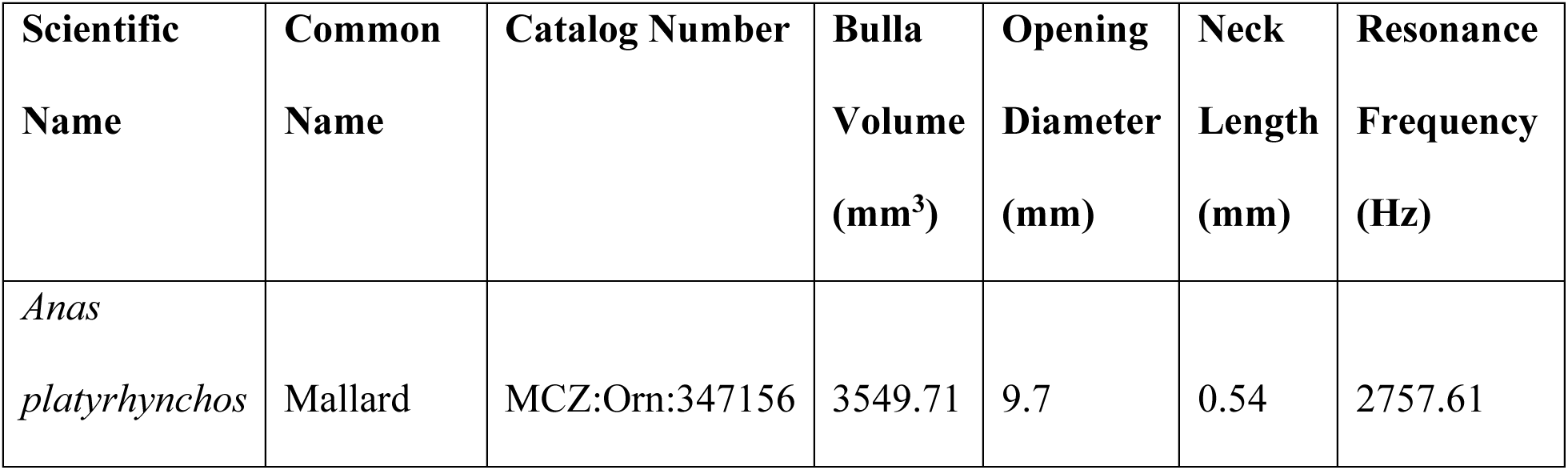
Measurements from µCT scan of mallard syrinx.

| Scientific<br>Name | Common<br>Name | Catalog Number | Bulla<br>Volume<br>(mm<sup>3</sup>) | Opening<br>Diameter<br>(mm) | Neck<br>Length<br>(mm) | Resonance<br>Frequency<br>(Hz) |
| --- | --- | --- | --- | --- | --- | --- |
| *Anas*<br>*platyrhynchos* | Mallard | MCZ:Orn:347156 | 3549.71 | 9.7 | 0.54 | 2757.61 |

**Table 2.**
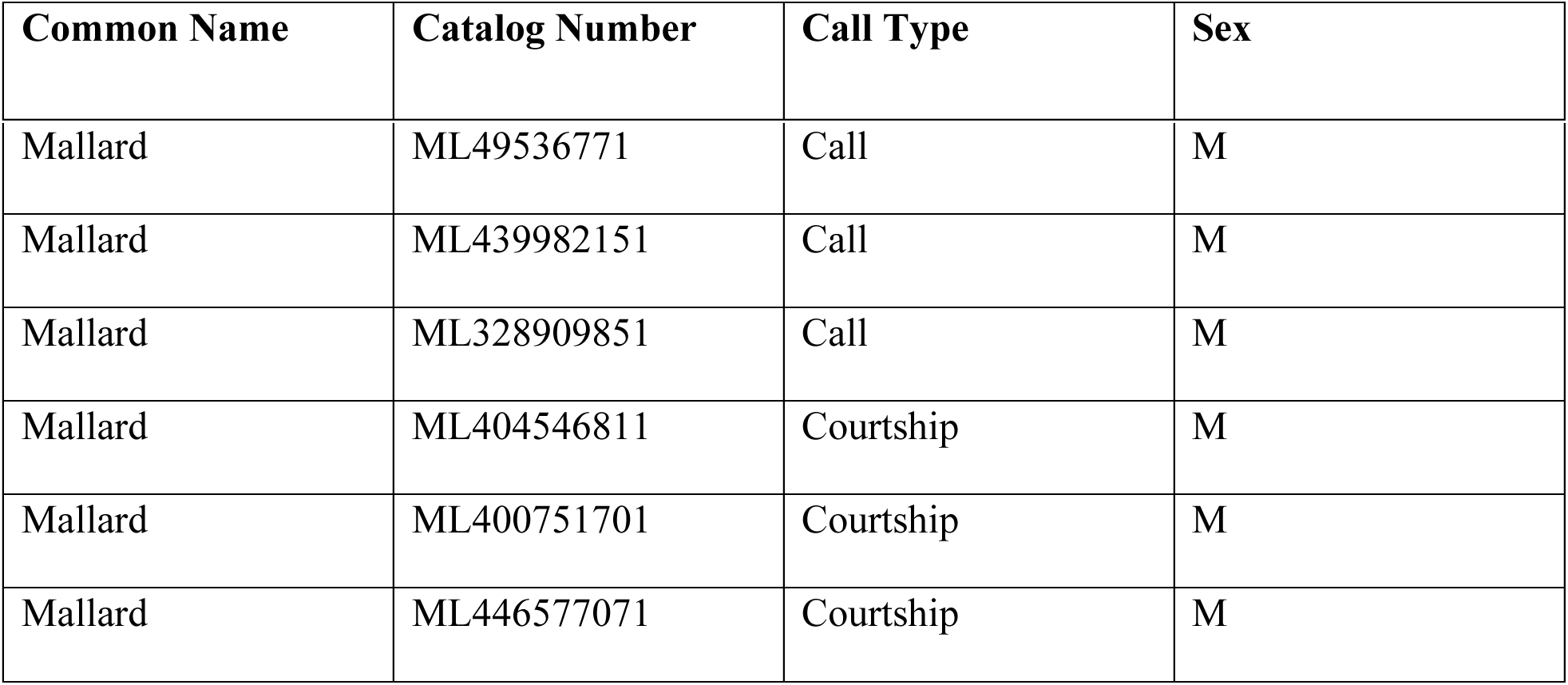
Sound files used to generate power spectra.

Evidence of an adaptive significance for the bulla leads to the question of how this left-right asymmetric, sexually dimorphic structure is produced. To address this, we first focused on the mechanism underlying its asymmetric formation. PITX2 is a transcription factor, induced in the left lateral plate mesoderm by NODAL, which has been shown to act upstream of left-right asymmetric morphogenesis of internal organs, including the heart, lungs and gut, in both mouse and chick^13,17,18,20,22^. As expected, when *PITX2* expression was examined by *in situ* hybridization chain reaction (HCR), strong *PITX2* expression was observed in the left side of the Hamburger Hamilton (HH) stage 13 duck splanchnic mesoderm, the lateral plate derivative that is the precursor to the mesoderm of the gut and airway, including the syrinx (Figure 2a)^23^. This early *PITX2* activity declines over subsequent stages, with little to no asymmetric *PITX2* expression being detected by HH17. *PITX2* expression is then reestablished (a second “wave” of expression) in the left half of the syrinx anlage by HH18, just prior to when the syrinx separates from the gut and forms an individual tube. In contrast, after a similar early level of asymmetric expression in the chick splanchnic mesoderm, the expression of *PITX2* undergoes a steady decline, does not rebound, and does not show asymmetric expression in the forming chick syrinx, which is a bilaterally symmetric structure in this species (Figures 2b, S2).

**Figure 2.**
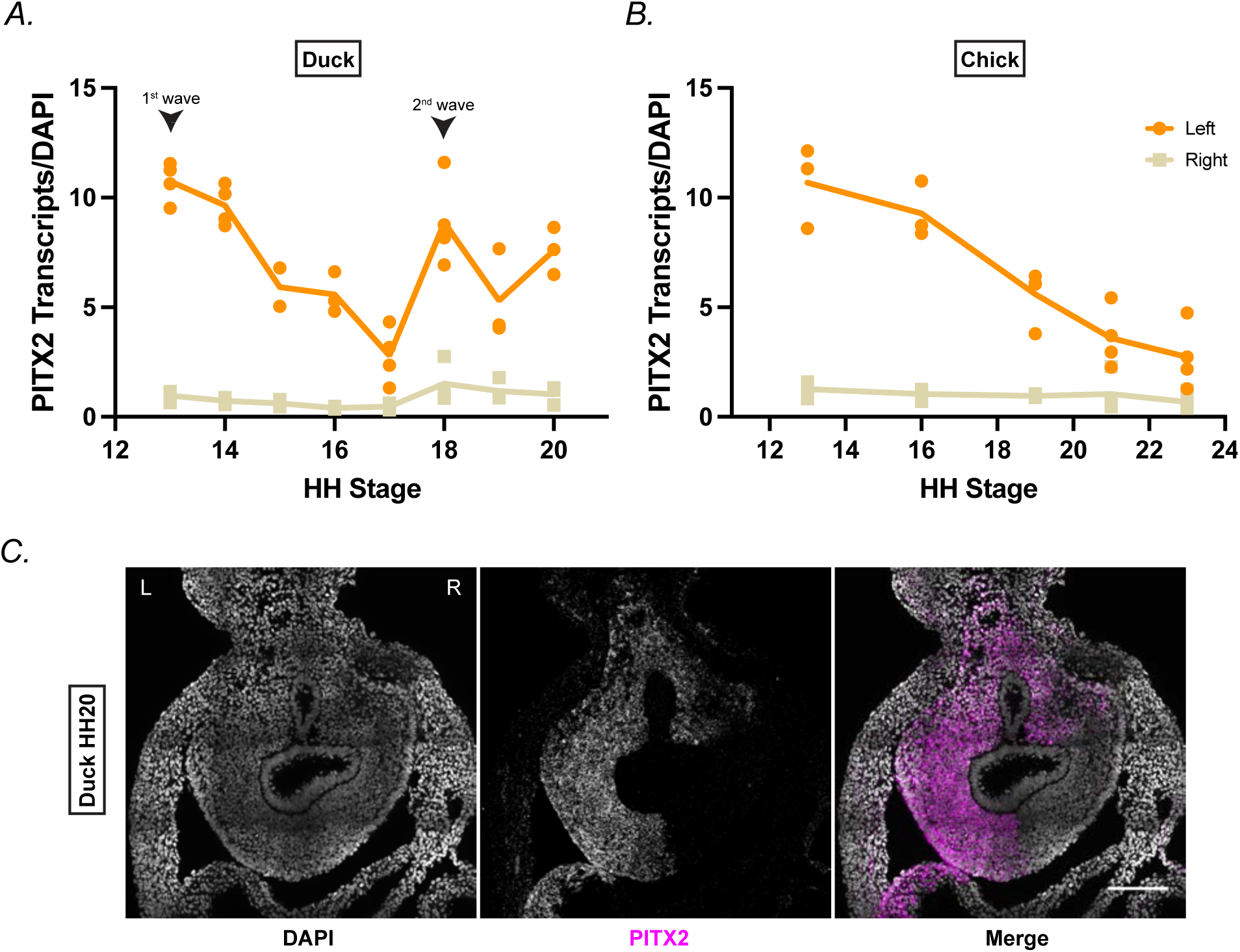
PITX2 shows two waves of expression in the splanchnic mesoderm and foregut in duck, not chick. PITX2 expression on the left vs. right in duck (A) and chick (B) in lateral plate mesoderm and ventral tracheal-esophageal tube. Measurements are an average of the PITX2 counts from 4 regions of interest divided by the DAPI counts per half. Each point represents a replicate with the line connecting time points at the mean. Probes generated against all PITX2 isoforms; PITX2a and PITX2b expected bilaterally while PITX2c is expected to be asymmetric. Duck early to late HH stage n = 4,4,2,3,4,4,3,3. Chick early to late HH stage n = 3,3,4,4,4. (C) Maximum intensity projection of HH20 duck airway. Right (R) and left (L) as labeled on image. Scale bar = 100µm.

While the first wave of expression of *PITX2* in the lateral plate is known to be induced by NODAL signaling, NODAL is absent by the time *PITX2* expression reinitiates in the duck airway^15,21^. A similar phenomenon has been described in the developing chick gut. After a decline in an initial asymmetric expression pattern. The chick midgut mesentery experiences a second wave of asymmetric *PITX2* activity. In the midgut, the trigger for the second wave was found to be TGFβ^21^. However, when we examined pSMAD2,3 immunostaining—a marker of active TGFß signaling—in the forming duck syrinx, we observed no expression at the syringeal level at the time of *PITX2* reactivation (n=3, data not shown). In contrast, pSMAD1,5,9, a marker of BMP signaling, shows immunostaining at the syringeal level coincident with the second wave of *PITX2* expression (Figure 3a), suggesting that BMP ligands could be at least partially responsible for *PITX2* reactivation. We also observe that BMP signaling is active in the chick over a similar time course (Figure S3).

**Figure 3.**
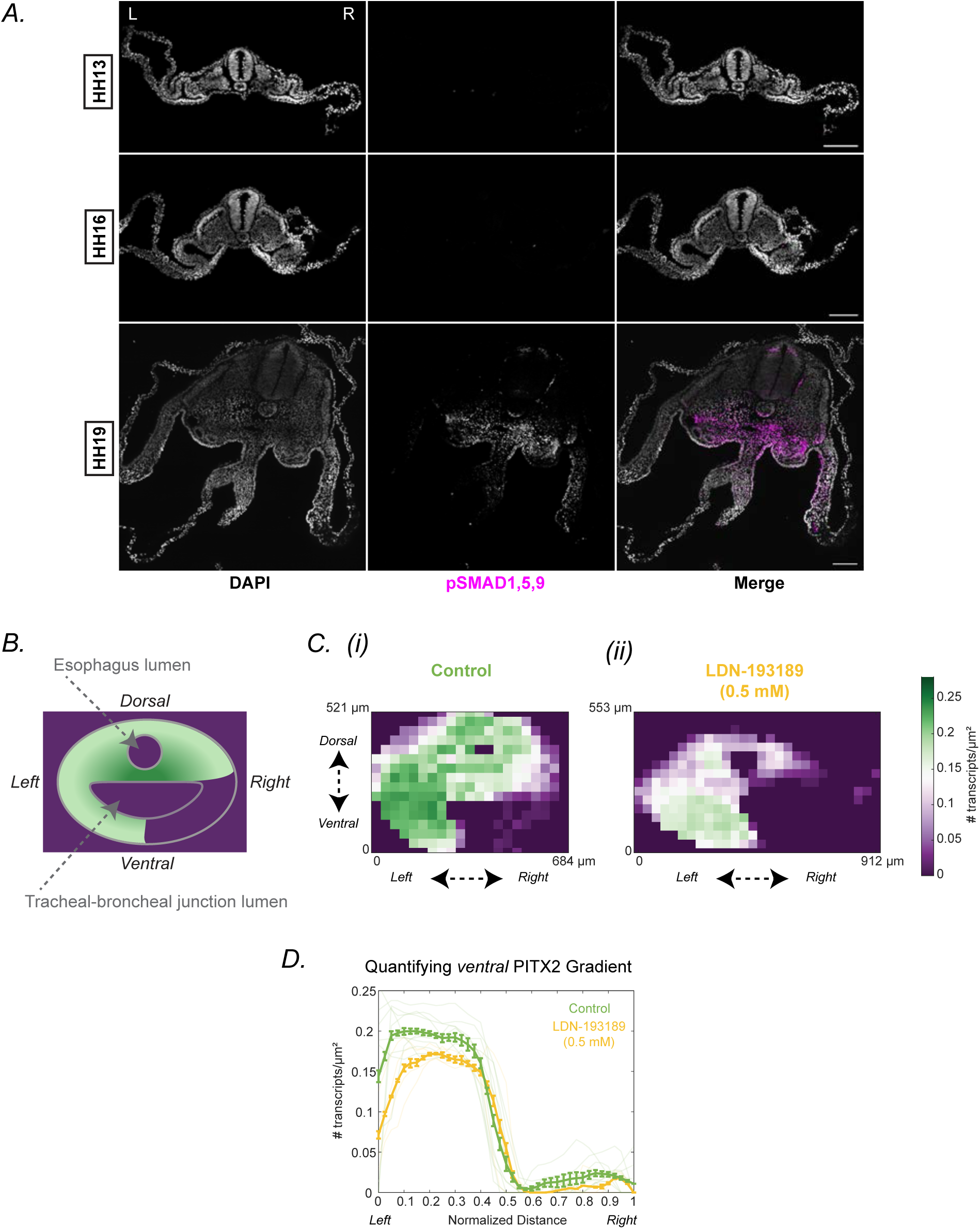
BMP is expressed at the time of the second wave of PITX2 and its inhibition results in decreased expression of PITX2. (A) Maximum intensity projections of representative HH13 (4/4), HH16 (3/3), and HH19 (3/3) duck embryos stained with pSMAD1,5,9. Right (R) and left (L) as labeled on image. Scale bar = 100µm. (B) Cartoon of transverse section of the tracheal-esophageal tube at the level of the tracheal-bronchial junction at HH21. Level of PITX2 transcripts represented from green (high) to purple (low). (C) Heat map showing PITX2 HCR of control (i) and LDN-193189 (0.5mM, BMP antagonist) -treated (ii) sections. (D) Quantification of the ventral (syrinx-fated) region of the heat maps shows significant decrease in transcript count in the treated group (yellow) vs. the control (green). Semi-transparent lines show individual samples. Line with error bars shows mean ± SEM.

To identify members of the BMP family that could be responsible for the observed pSMAD1,5,9 activity we conducted a screen for BMP genes expressed at the period when PITX2 is upregulated for the second time. We found that two BMP family members, BMP4 and BMP5, exhibit high levels of expression at this later stage (Figure S9). To test the possibility that BMP signaling might be involved in regulating the second wave of *PITX2* expression in the anlage of the duck syrinx, we used the BMP inhibitor LDN-193189 dihydrochloride (an inhibitor of ALK2 and ALK3). After 1 day of *in ovo* treatment, airways showed diminished (although not eliminated) pSMAD1,5,9 expression at HH21 (Figure S4). BMP inhibition resulted in an altered domain of *PITX2* expression and significantly reduced levels of *PITX2* RNA in the ventral region, which will develop into the syrinx (Figures 3b–d, S5). Thus, BMP activity appears to contribute to the reactivation of *PITX2* expression in the second wave.

These results indicate that the second wave of expression of *PITX2* on the left side of the duck airway is induced by BMP signaling, just as the first wave of *PITX2* expression in the left LPM is induced by NODAL. Initially, *PITX2* is expressed exclusively on the left during its initial wave of expression due to asymmetric NODAL. However, BMP signaling appears to be bilaterally symmetric at the time of the second wave of *PITX2* expression, thus raising the question of how BMP activity is interpreted asymmetrically. An attractive hypothesis is that this could be encoded as a residual consequence of the first wave of *PITX2* expression in the form of epigenetic memory. Indeed, creating such a positional memory throughout the left LPM might be the reason the first wave of *PITX2* expression takes place, prior to any asymmetric morphogenesis. To explore this question, we conducted a single-nucleus multiomic study, including both RNA-sequencing (RNA-seq) and assay for transposase-accessible chromatin with sequencing (ATAC-seq), on separately collected left and right-side tissue fated to form the syrinx, harvested from stage 13, 16-17, 21, and 25 duck embryos. The RNA-seq data in this analysis confirmed our earlier observation that *PITX2* is strongly expressed in the stage 13 left LPM (Wave 1), and that this expression is lost by stage 16–17. Subsequently, *PITX2* expression increases again on the left side through stages 21 and 25 (Figure 4a). Not unexpectedly, the ATAC-seq data indicates that the chromatin at the *PITX2* promoter is open on left but not the right side of the LPM at stage 13, but in striking contrast to the RNAseq data, this asymmetric accessibility is maintained at all later stages assessed (Figure 4b).

**Figure 4.**
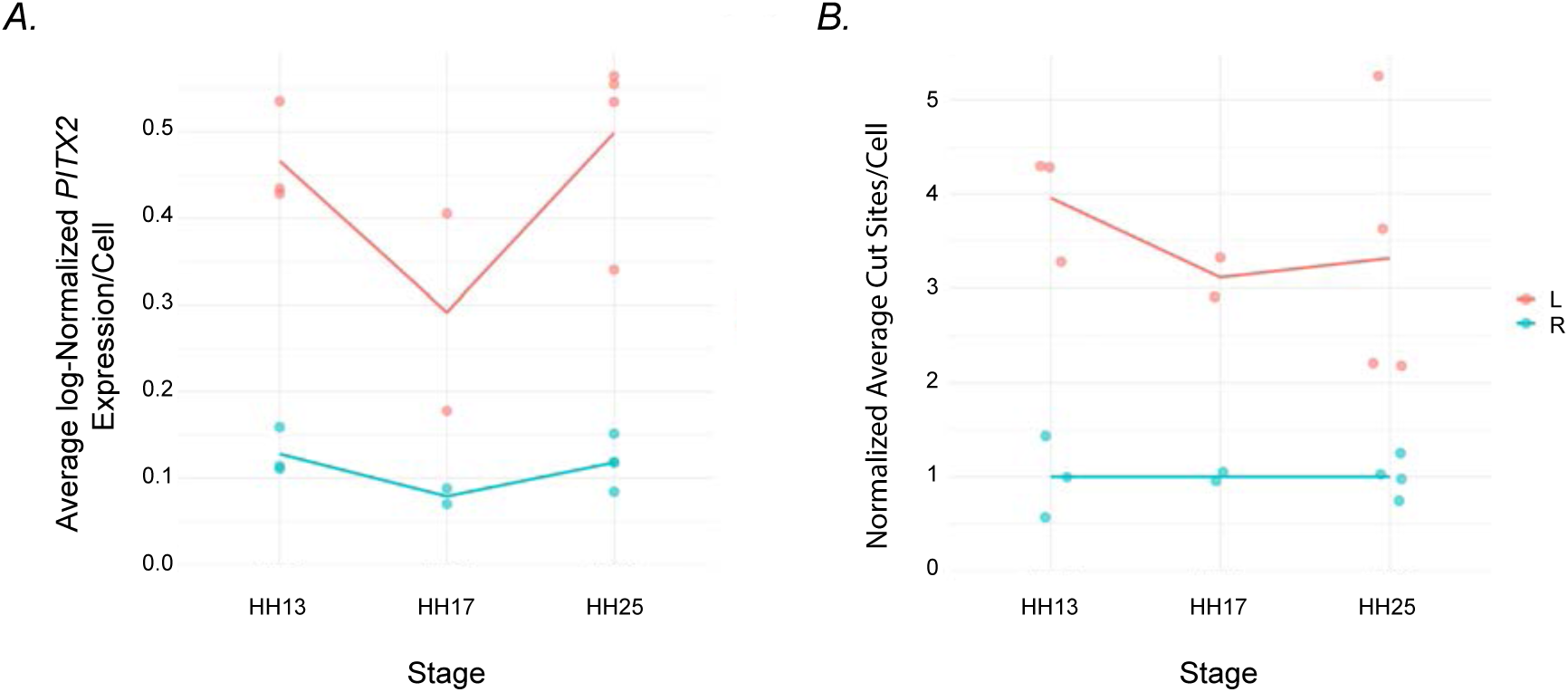
Single-nucleus RNA-seq and ATAC-seq show left-sided RNA expression in two waves and stably accessible left-sided chromatin. (A) Single-nucleus RNA-seq shows average normalized *PITX2* expression on the right (blue) is low from HH13–HH25. Average normalized *PITX2* expression on the left (red) is elevated at HH13 (wave 1), decreased at HH17, and then elevated again at HH25 (wave 2). (B) Single-nucleus ATAC-seq shows elevated levels of chromatin accessibility to the left from HH13– HH25.

Having established that left-right asymmetric positional information is present in the forming duck syrinx, we next wanted to know the cellular properties being asymmetrically regulated in the duck syrinx to produce a left-specific bulla, and the asymmetric gene activity responsible for driving them. To this end, we first established a time-course of male and female syrinx development. The syrinx initially forms as a single symmetric structure with no visible left-right difference or sexual dimorphism. By embryonic day 10 (E10), a small asymmetric growth can be observed on the left side of both female and male duck syrinxes. However, two days later this asymmetry can no longer be seen in the female specimens and the female syrinx continues to grow symmetrically. In contrast, the left-sided outgrowth in the male duck syrinx continues to expand on the left side at E12, and subsequent stages (Figure 1b).

To address the cellular processes responsible for this growth, syrinxes were collected at various stages and assessed for the level of cell death, cell number, and proliferation. From these experiments we concluded that differential growth of the bulla is due to asymmetric levels of cell division and not because of changing cell density or levels of cell death (Figure 5a). Data revealed virtually no cell death in the syrinx from E6 to E10 (<0.6%), as measured through immunofluorescence with a cleaved caspase-3 antibody (data not shown). Similarly, we did not observe any significant differences in density across the samples, measured as number of nuclei per μm^2^. In contrast, proliferation, assessed by incorporation of the thymidine analogue EdU shows higher proliferation in the left side of male duck syrinxes at E9 and E11 (Figure 5a).

**Figure 5.**
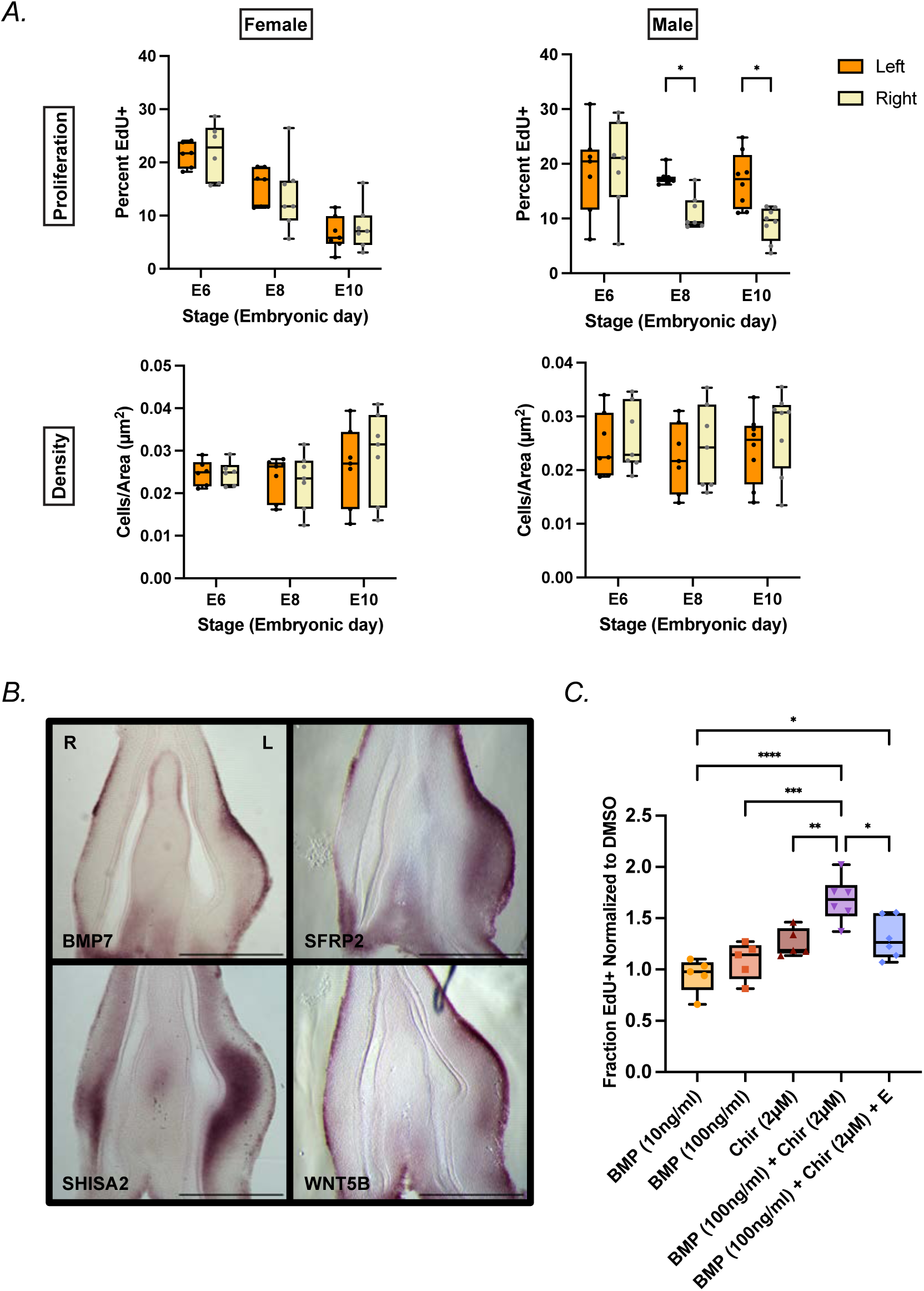
Proliferation is responsible for asymmetric syringeal morphogenesis and the BMP and WNT pathways influence proliferation. (A) Cell proliferation, not density, is higher on the left than on the right in males. Female and male syrinxes were treated with a 1 hour EdU pulse before being sacrificed. Single sections were evaluated for total cell number, EdU^+^ cell number, cleaved caspase-3 stain, and total area. Plots show female and male samples from embryonic day 6 to embryonic day 10 (E6 to E10). Left vs. right tested with Wilcoxon matched pairs signed rank test with Holm-Šídák post hoc. *p<0.05. Cell death not shown, but never exceeded 0.6% of cells. n = 7 except FE6, n = 6, ME10, n = 8. (B) *In situ* hybridizations of gene candidates generated from the scRNA-seq dataset. All samples are E9 male. E9 female samples show same patterns of expression. Right (R) and left (L) as labeled on image. Scale bar = 0.5mm. (C) Proliferation rates in E7 micromass cultures treated with BMP and WNT agonists with respect to DMSO control. Data points are the average of two technical replicates. BMP4 was assayed at 10 ng/ml and 100 ng/ml, CHIR99021 (Chir; WNT agonist) at 2µM, and both were assayed in combination with Chir 2 µM and BMP4 100 ng/ml. The Chir/BMP4 cultures were then assayed with 100 nM estradiol (E). n (left to right) = 5,5,5,6,6. 1-way ANOVA with Holm-Šídák post hoc. *p<0.05, **p<0.005, ***p<0.0005, ****p<0.0001.

Having established that the growth of the left-sided bulla is driven by higher proliferation rates, we next looked for pathways that might be asymmetrically regulated in the syrinx to modulate levels of cell division. To that end, we conducted single cell RNA-sequencing (scRNAseq) analysis on syrinxes from E6 (when the syrinx is symmetric), E9 (when slight left-biased asymmetric growth is present in both male and female), and E11 (when the asymmetry of growth is male-specific) (Figure S6). We focused on genes expressed in mesodermal cell clusters in our scRNAseq analysis. A number of genes implicated in this way were components of the WNT signaling pathway (Figure S7). In addition, our attention was drawn to BMP signaling (Figures S7, S8). Candidates were screened by *in situ* hybridization. This led to the identification of a number of genes, including BMP7, SHISA2, SFRP2, and WNT5b, with strikingly asymmetric left sided expression in the developing syrinx (Figures 5b, S10). These expression patterns suggest that WNT and BMP signaling might play key roles in duck syrinx morphogenesis.

To test whether left-sided BMP and WNT activity were indeed involved in promoting proliferation in the syrinx airway primordium, we turned to an *in vitro* micromass culture system. Cells from E7 syrinxes were dissected, dissociated, and plated in 24-well plates (2 cm^2^ per well) at 9,000 cells/µl. After allowing the cultures to establish for 24 hours, micromass cultures were treated for a further 24 hours before collection. We observed a trend towards higher proliferation rate with the addition of either BMP4 protein or WNT agonist (CHIR99021). Moreover, in combination, these factors synergistically and significantly increased the proliferation rate of cells as compared to DMSO-treated control cells (Figure 5c).

Together, these data suggest a model where asymmetric activation of *BMP* and *WNT* activity drives left-specific growth of the bulla in the duck syrinx. However, as BMP and WNT pathway genes are initiated equivalently in developing male and female duck syrinxes (Figures 5b, S10), the question is raised of how the sexual dimorphism of bulla formation is controlled. We hypothesized that this must be tied to the sex steroids that differentially direct male (androgens) and female (estrogen) characteristics. Moreover, results from castration experiments show that the absence of gonads and hence loss of sex steroid production produces a male-like syrinx with a left-sided bulla in males and females^6^. This implies that although the left-right pathway is capable of driving bulla formation in both sexes, it is prevented from doing so by the activity of estrogen in females. Consistent with this, at E10, prior to estrogen production in the female, increased left-sided syringeal growth can be observed in both males and females. However, by E12, after gonad maturation at E9–11, asymmetric growth of the bulla is exclusively seen in males (Figure 1b)^3,6,24^. To see whether estrogen signaling might be acting directly in the proliferating cells forming the left-sided bulla, we examined expression of estrogen receptors. We surveyed all estrogen receptors that appeared in our E9 scRNAseq dataset and of the six genes examined, only ESR1 is widely expressed in the mesodermal cells in the syrinx (Figure 6a). Moreover, it, too, is left-specific in its expression. Notably, its expression was found in both male and female samples. *In situ* hybridization for ESR1 confirms left-sided expression in males and females at E9 and E11 (Figure 6b, S10).

**Figure 6.**
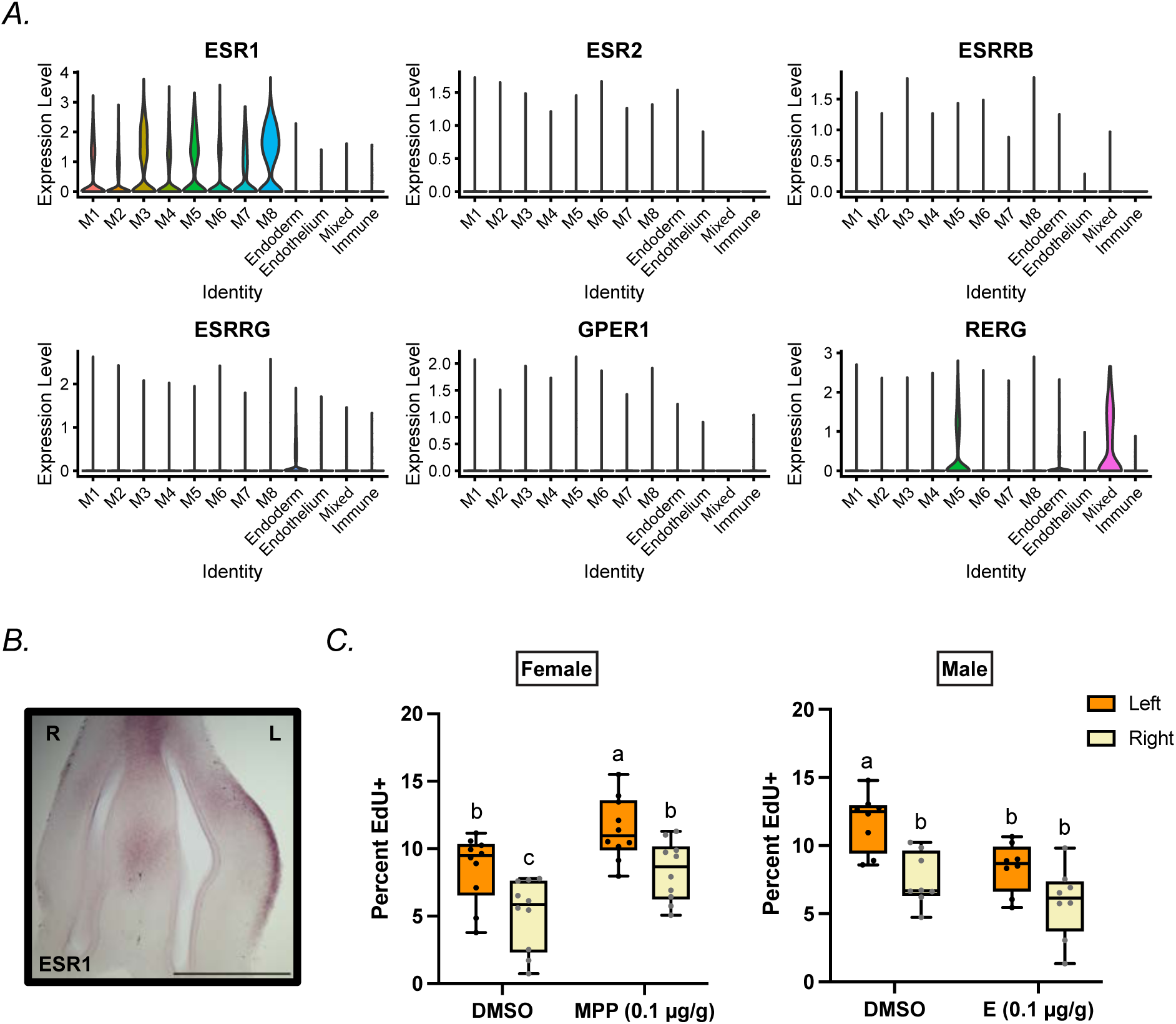
ESR1 shows left-sided expression in the syrinx and is upstream of cell proliferation rates. (A) Violin plots of all expressed estrogen receptors in the data set. Clusters M1–M8 are mesoderm clusters 1–8. Mixed refers to a population including fibroblasts, smooth muscle, and immune cells. ESR1 is the only receptor to show expression throughout the mesoderm. (B) *In situ* hybridization of E9 male syrinx for ESR1 shows left-sided expression. Right (R) and left (L) as labeled on image. Scale bar = 0.5 mm. (C) Percent EdU^+^ cells in syrinxes from females and males treated *in vivo* with MPP dihydrochloride (a specific ESR1 inhibitor) and estradiol, respectively. Treatment administered at E7 and tissue collected after 1 day. 1-way ANOVA with Tukey’s post hoc. Female, n = 10. DMSO_L vs. DMSO_R: p = 0.0174, DMSO_L vs. MPP_L: p = 0.0496, DMSO_L vs. MPP_R: p = 0.9975, DMSO_R vs. MPP_L: p < 0.0001, DMSO_R vs. MPP_R: p = 0.0279, MPP_L vs. MPP_R: p = 0.0317. Male, n = 8. DMSO_L vs. DMSO_R: p = 0.0025, DMSO_L vs. E_L: p = 0.0201, DMSO_L vs. E_R: p < 0.0001, DMSO_R vs. E_L: p = 0.8377, DMSO_R vs. E_R: p = 0.4310, E_L vs. E_R: p = 0.1067.

The expression of *ESR1* in males should not alter syringeal growth, as no estrogen is present to activate the receptor. However, signaling through ESR1 In females could act to oppose the proliferative effect of left-sided WNT and BMP activity. To test his hypothesis, we returned to the micromass assay. When we added estradiol to our micromass cultures containing WNT and BMP agonists, we indeed saw a decrease in proliferation rate (Figure 5c). We additionally tested the effect of perturbing estrogen signaling *in ovo.* We treated E7 females with MPP dihydrochloride, an ESR1-specific antagonist. After one day of treatment, EdU staining showed an increase in proliferation rates. Conversely, males treated in an analogous manner with estradiol showed a decrease in proliferation. Interestingly, in the males treated with estradiol, only the left side, where the ESR1 receptor is present, showed a significant change in proliferation rate (Figure 6c).

## Discussion

Intense intra- and intersexual selection in ducks has led to the evolution of sexual dimorphism in several traits, likely including the male-specific left-sided syringeal bulla ^7,25,26^. The species-specific plumage and courtship displays of male ducks may act as premating barriers to gene flow, especially in contexts where many different species coexist as is true for mallards^27,28^. We show evidence that the mallard bulla contributes to the production of male, sexually dimorphic vocalizations used during courtship displays through its resonance properties. While further work is needed to test the direct connection between the bulla and sound production, this result suggests its function and that the bulla is a sexually selected trait.

We show that spatiotemporally dynamic *PITX2* expression is essential for duck syrinx organogenesis. PITX2 is known to play a central role in establishing early left-right asymmetry and later in organ laterality^15,29^. Much work has gone into the early determinants of establishing left-right asymmetry, but work on organ-specific laterally asymmetric development is more limited^19,20,22,30–32^. We observe that after the first wave of left LPM-specific PITX2 expression abates, a second is induced—likely by bilateral BMP signaling—as a left-sided syrinx-specific signal (Figure 7a). In contrast, Sanketi et al. (2022) found that TGFß induces a similar second wave to induce left-right asymmetry in the chick gut, while we see no evidence of active TGFß signal in the syrinx primordium. Thus, while the overall PITX2 expression pattern is conserved, different organs show distinct regulatory triggers for its expression.

**Figure 7.**
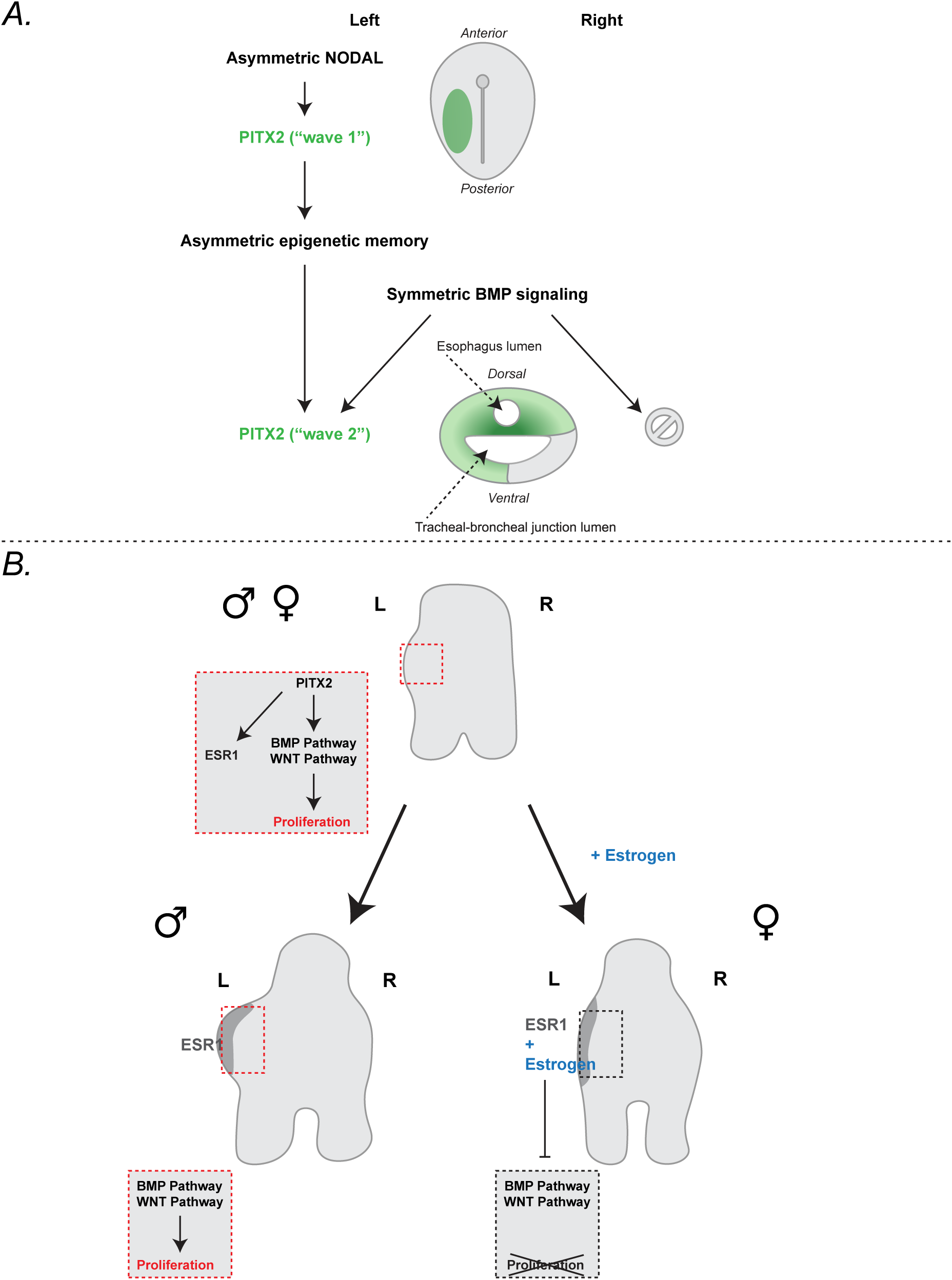
Models of the two waves of PITX2 and integration of left-right asymmetry with sexually dimorphic developmental plans. (A) Early left-sided NODAL expression activates a first, left-sided wave of PITX2 expression, which declines by HH17. Maintenance of asymmetric chromatin accessibility allows bilaterally expressed BMP to activate a second wave of left-sided PITX2 expression at HH18. (B) Left-sided PITX2 is upstream of left-sided ESR1 and BMP/WNT pathways. BMP/WNT pathways are in turn upstream of elevated proliferation. In females, as estrogen is produced it binds ESR1, inhibiting elevated proliferation.

Using a single-nucleus multiome approach we found that a major function of the initial, LPM-specific asymmetry may be to set up an epigenetic memory for asymmetrically developing tissues. We observed, as in the chick gut, that the initial PITX2 asymmetry from the left LPM was carried into the left-splanchnic mesoderm but then subsided before returning in the forming syrinx. This second wave is induced by a bilateral signal, and thus is dependent on differentially accessible chromatin to induce left-sided expression. Establishment of chromatin memory has been shown to play a role in processes such as in the inflammatory response. In the inflammatory response, transcription factors bind specific memory domains, resulting in poised, open chromatin. This allows for a rapid response that can be activated by diverse stimuli other than those that initially triggered the response^33^. If a similar mechanism is involved in the avian left-LPM, this may explain how the chick and duck show related but diverged regulation in the gut and the syrinx.

The establishment of left-right positioned information in the developing duck syrinx leads to left-specific activation of BMP and WNT signaling, which in turn drives left-sided proliferation and hence asymmetric development of the syringeal bulla (Figure 7b). This left-sided cascade, and hence the impetus for bulla formation, occurs in the syrinx primordium of both male and female ducks, and as a consequence there is an initial left-sided outgrowth on the developing syrinx of both sexes. However, once embryonic sex steroid production is initiated, this increased left-sided proliferation is suppressed in females. To this end, in addition to pro-proliferative signals, the estrogen receptor ESR1 is upregulated on the left side of both the male and the female syrinx. In the male this makes no difference, as there is no estrogen to activate the receptor. However, in females, activation of left-localized ESR1 by estrogen results in inhibition of proliferation and therefore of asymmetric syringeal development.

The confluence of the left-right cascade and sex-hormone signaling is uncommon but also exists in the development of avian gonads^34–37^. With only a few exceptions, male birds develop symmetric gonads while females develop asymmetrically with maturation only occurring on the left^38^. While the eventual syrinx asymmetry is in the male, not the female, many aspects of gonadal development are reflected in the syringeal developmental plan. Work in the chick gonads has implicated the NODAL-PITX2 cascade^39^. Further, both males and females show early left-sided expression of ESR1 in the gonadal epithelium and manipulations have shown that PITX2 is upstream of *ESR1* expression in this system^34,35^. Other intriguing connections include that BMP7 has been observed to have left-sided expression in both sexes in the indifferent ridges that develop into chick gonads and expression of PITX2 on the right induces right-sided expression in the chick gonad^34,40^. We observe asymmetric ESR1 and BMP7 in the duck syrinx in both males and females, indicating overlap in the gene regulatory networks involved in both systems.

Taken together, our results suggest that the asymmetric, male-specific bulla in the duck syrinx evolved under sexual selection to produce a unique mating call. To accomplish this, evolution took advantage of early asymmetric changes in the developmental chromatin landscape to establish the bulla on the left and harnessed the sex-specific production of estrogens to assure that the structure is only produced in males. This work thus highlights how distinct signaling systems can be integrated during embryonic morphogenesis to achieve anatomical adaptation.

## Methods

### Egg sourcing and incubation

Two breeds of duck, Mallard and Khaki Campbell, were used throughout this study. They were considered interchangeable unless otherwise specified. Eggs were sourced from Metzer Farms (Monterey County, CA) and from Chad Araneo (MA). Chick eggs were Specific Pathogen Free (SPF) White Leghorn chicken eggs from Charles River Laboratories and AVS Bio. Roslin Green cytoplasmic GFP chicken eggs^41^ were sourced from Susan Chapman (Clemson University Biological Sciences, Clemson, SC). All eggs were incubated at 37°C at 45% humidity. Duck and chick embryos were staged using the Hamburger and Hamilton (HH) staging series^23^. Up to HH25 (E6) embryos are staged using the HH series, while for later time points embryonic day is used.

### DNA extraction and embryo sexing

At E2-3 eggs were lowered by removing 5ml of albumin. At E5–6 eggs were windowed and for each egg a small piece of the amnion was removed and placed in 100μl of 50mM NaOH. Samples were incubated for 1 hour at 95 C° after which 25μl of 1M Tris pH8 was added to each sample. 1μl of the DNA extraction solution was used for PCR; conditions and primer sequences as in Li et al., 2015^42^.

### Alcian Blue stain

Airway tissues were dissected in PBS and fixed overnight in PBS-buffered 4% formaldehyde at 4°C, then washed 2x in PBS before staining overnight at room temperature in Alcian Blue stain (0.02% [w/v] Alcian Blue 8GX, 5% [v/v] glacial acetic acid in 70% ethanol). This was followed by 3x 30min washes in 95% ethanol and 1 hour in 1% KOH. Samples were destained in fresh 1% KOH, repeating until tissues no longer leeched stain. Samples were stored at room temperature after being graded through 20% glycerol/0.8% KOH, 40% glycerol/0.6% KOH, 60% glycerol/0.4%KOH, 80% glycerol/water on a rocking platform.

### Cryosectioning

For pre- and post-hybridization HCR, immunostaining, and EdU staining, (described below) formaldehyde-fixed tissues were graded through 10% Sucrose in PBS (w/v), 30% Sucrose in PBS (w/v), and 1:1 (v/v) 30% Sucrose in PBS: OCT (Tissue-Tek^®^ O.C.T. Compound, 4583). Samples were embedded in OCT and stored at -80°C. Embedded airways were cryosectioned onto Superfrost^TM^ Plus microscope slides (Fisherbrand, 12-550-15) at 16–18µm thickness.

### *In situ* hybridization

Whole mount *in situ* hybridizations done as described in Longtine *et al*., 2024 with the following exception^43^. Probes were visualized with NBT/BCIP (Roche 11681451001). Samples embedded in albumin/gelatin (10% sucrose w/v, 30% BSA w/v, 0.5% gelatin w/v in PBS). Samples equilibrated in albumin/gelatin, then embedded in albumin/gelatin media mixed with glutaraldehyde (Sigma-Aldrich G7651, 100µl 12.5% glutaraldehyde combined with 1ml embedding media). Sectioned at 50 µm on the vibratome (Leica VT1000M 046227584) and imaged as described below.

### Hybridization chain reaction (HCR) *in situ* hybridization

HCR probe sets were designed using HCR probe maker 3.7 V2 from the Özpolat lab (https://github.com/rwnull/insitu_probe_generator/tree/master)^44^. Probes were synthesized by Integrated DNA Technologies and individually resuspended to 10nM with H_2_O. All probes were then pooled for a final concentration of 10µM. and stored at -20°C. Sequences for the duck *PITX2* and chick *PITX2* probe sets are in **Error! Reference source not found.**.

Samples were dissected in PBS and fixed for one hour in PBS-buffered 4% formaldehyde. HCR *in situ* hybridization was conducted as described in Choi *et al*., 2018, with exceptions outlined in Choi *et al*., 2024^45,46^.

After HCR, whole airways were embedded and cryosectioned as described above. Sections were washed for 5 minutes in PBS before incubation for 40 minutes with 300 nM DAPI in PBS. Slides were washed 2x 5 minutes with PBS and cover slipped with Fluoromount-G (Southern Biotech, 0100-01) before being stored at -80°C until imaging.

### Section HCR

For HCR on previously cryosectioned airways, tissues were dissected, fixed, embedded, and cryosectioned as described above. HCR was performed using a modified version of the whole-mount protocol, which differs as follows.

Slides were removed from -80°C storage, washed with PBS for 5 minutes at room temperature. After the PBS was removed, the slides were positioned vertically and the tissue sections air dried for 5 minutes. Sections were washed with PBS for 5 minutes before permeabilizing with 70% ethanol/30% PBS (v/v) for 5 minutes, then washed 2x 5 minutes in 1x PBS. The following incubations all took place in a humidified chamber. Slides were incubated in a preheated chamber with preheated hybridization buffer (Molecular Instruments) at 37°C for 10 minutes. The probe mixture was prepared as for whole-mount HCR and sections were incubated overnight in the probe solution at 37°C, covered with a plastic coverslip cut from 4mm LDPE bag (VWR, 89072-302). Coverslips were removed and sections were washed 2x 30 minutes at room temperature in wash buffer (Molecular Instruments) preheated to 37°C. Sections were then washed 2x 20 minutes with 5x SSCT at room temperature, followed by a preincubation in amplification buffer (Molecular Instruments) for 30 minutes at room temperature. Hairpins were prepared as for whole mount HCR, and sections were incubated overnight at room temperature in the hairpin solution under plastic coverslips, protected from light. Slides were washed 2x 20 minutes in 5x SSCT at room temperature, followed by DAPI staining and mounting as described above.

### HCR image quantification for chick vs. duck comparisons

Individual RNA transcripts were counted using the Matlab (Mathworks) plugin ImageM^47^. *PITX2* expression on the left vs. right in duck and chick are the average of the *PITX2* counts from 4 regions of interest per side divided by the DAPI counts.

### Image quantification for ventral *PITX2* expression

#### Image Processing and Spot Detection

HCR images were processed using ImageJ (Fiji), where they were aligned, cropped, and analyzed for RNA spots using TrackMate with a spot diameter of 3 pixels. The centroid coordinates and intensity data of the detected spots were recorded and exported for further analysis in MATLAB. Mask files, defining regions of interest (ROIs) and excluding unwanted areas such as the lumen or background, were created and imported as TIFFs into MATLAB.

#### Transcript Spot Quantification

For each mask, a binary image was generated based on the centroid coordinates of the detected spots. The corresponding mask was applied to restrict the analysis to the ROI. Transcript spot counts were computed using the *blockproc* function in MATLAB, dividing the image into 100×100-pixel blocks. Blocks with counts below a threshold of 12 were excluded to avoid erroneous edge effects, which typically result from insufficient data or partial block coverage at the boundaries. The counts were then normalized by the area of each block (in µm²) and visualized as spatial maps, with a color scale indicating transcript density.

#### Region Segmentation and Normalization

Ventral regions were defined based on user-specified boundaries for each sample. The mean transcript density and standard error of the mean (SEM) were calculated for each region. To enable comparison across samples of varying sizes, the horizontal distance was normalized, with 0 corresponding to the leftmost pixel and 1 to the rightmost. Additionally, the normalized distance was interpolated to generate 40 data points between 0 and 1 (with a step size of 0.025) to ensure consistent comparison across samples.

#### Statistical Analysis and Visualization

Transcript densities for the ventral regions were plotted along a normalized left-right distance axis with error bars representing the SEM. Two-sided Wilcoxon rank sum test used to test for significance.

### Immunostaining

#### Immunostaining of sections

Cryosections (obtained as described above) were washed 2x 5 minutes in PBTS (0.2% [w/v] BSA [Sigma-Aldrich, A9647], 0.1% [v/v] Triton X-100 [Sigma-Aldrich T8787], 0.02% [w/v] SDS [Sigma-Aldrich, 11667289001] in PBS). Sections were blocked for 30 minutes at room temperature in PBTS + 10% serum matched to the secondary antibody species, followed by overnight incubation at 4°C with primary antibody in PBTS (dilutions as indicated below). Slides were then washed 2x 10 minutes in PBTS and incubated with secondary antibody in PBS (1:500 dilution, donkey anti-rabbit [Jackson ImmunoResearch, 711-605-152 or 711-165-152]) for 1 hour at room temperature, followed by 2x 10 minutes washes in PBTS and a final 5 minute PBS wash. Tissue sections were counterstained with DAPI and mounted (as described above) for imaging.

#### Immunostaining of sections with antigen retrieval

For phospho-SMAD staining, antigen retrieval was performed with citrate buffer at pH 6.0. Cryosections were washed 2x 5 minutes in PBS, followed by 3x 5 minutes in TN buffer (0.1M Tris-HCl, pH 7.5). Slides were placed in a preheated slide mailer, covered with boiling citrate buffer, and placed in a vegetable steamer for 10 minutes. Slide mailers were then cooled on ice at 4°C for 15 minutes. Slides were then washed 2x 5 minutes in TN, washed for 5 minutes in TNT (TN with 0.05% [v/v] Tween20 [Sigma-Aldrich, 11332465001]), and blocked for 1 hour in TNTB (TNT with 0.5% blocking reagent [Roche Diagnostics Deutschland GmbH, 11096176001]) at room temperature. Sections were incubated with primary antibody (in TNTB; dilutions specified below) at 4°C in a humidified chamber overnight. Slides were then washed 3x 5 minutes in TNT and incubated for 2 hours with secondary antibody’s used above (1:300 in TNTB) and DAPI (300 nM) at room temperature, then washed 4x 5 minutes in TNT. Sections were mounted as described above for imaging.

#### Primary antibodies used

Cleaved Caspase-3 (Cell Signaling Technology Asp175 Cat. #9661S) at 1:400; Phospho-SMAD1 (Ser463/465)/SMAD5 (Ser463/465)/SMAD9 (Ser465/467) (Cell Signaling Technology D5B10 Cat. # 13820S) at 1:200; Phospho-SMAD2 (Ser465/467) (Cell Signaling Technology 138D4 Cat. # 3108S) at 1:200.

#### EdU labeling

Staining was conducted with the Click-iT™ EdU Cell Proliferation Kit for Imaging, Alexa Fluor™ 488 dye (Thermo Fisher C10337). ***In ovo***: 400µl 1mM EdU solution was pipetted onto embryos in windowed eggs and incubated at 37°C at 45% humidity for 1 hour. Embryos were then dissected, fixed, and cryosectioned as described above before beginning the manufacturer-provided immunofluorescence protocol. **Micromass**: For each well, half the micromass media was removed (250µl) and replaced with 0.02 mM EdU for 4 hours. Samples were fixed for one hour in 4% formaldehyde in PBS at room temperature and cells were washed 2x 5 minutes in 1X PBS before immediately beginning the manufacturer-provided immunofluorescence protocol.

#### Quantification

Immunostaining was quantified manually using FIJI to count total EdU+ cells, DAPI+ cells, and cleaved caspase-3+ cells in the central z-slices from imaged syrinx sections.

### Micromass cultures

Micromass cultures consisted of E7 left-sided dissociated syrinx cells seeded on 24-well culture plates (3.4 ml wells, Corning 3524) at 9,000 cells/µl (20µl per culture). 10–12 syrinx halves were pooled per experiment. Data points are the average of two technical replicates. Seeded cultures were incubated for 2 hours before adding 500µl media (DMEM [Thermo Fisher, 11995065]; 1% Pen Strep [Gibco 15-140-163]; 10% Fetal Bovine Serum [Gibco 10437028]). Cultures were established for 24 hours and then treated. Cultures were incubated with treatments for 24 hours before being stained with EdU (see EdU labeling, above). BMP4 (R&D Systems, 5020-BP) was assayed at 10 ng/ml and 100 ng/ml, WNT agonist CHIR 99021(Tocris, 4423) at 2µM, and both were assayed in combination with CHIR 2 µM and BMP4 100 ng/ml. The CHIR/BMP4 cultures were then assayed with 100 nM estradiol (Millipore Sigma, 50-28-2). Equivalent volumes of DMSO were used as controls.

### Imaging

Alcian blue stains were imaged with Leica M205 stereoscope with base-transmitted light. Colorimetric ISH sections were imaged on Nikon Eclipse E800. Fluorescent microscopy was conducted on spinning disk confocal Nikon Ti2 inverted microscope with W1 Yokogawa spinning disk with a 50µm pinhole disk. EdU and immunofluorescence were captured at 20x (Plan Apo λ 20x/0.8 DIC I) and HCR at 40x and 60x (Plan Fluor 40x/1.3 Oil DIC H/N2, Plan Apo λ 60x/1.4 Oil DIC).

### *In ovo* drug treatments

#### BMP inhibition

HH15 duck eggs lowered, windowed, and treated with 100µl of 0.375mM or 0.5 mM LDN-193189 dihydrochloride (Tocris 6053) or the same volume of PBS. Embryos collected at HH21 and fixed, embedded, cryosectioned, stained, and imaged as described above.

#### Estradiol and MPP cultur

Duck eggs sexed as described above. At E7 male eggs treated with 0.1µg/g estradiol (Millipore Sigma, 50-28-2), females with 0.1µg/g MPP dihydrochloride (Tocris 1991) diluted in 400µl PBS. Controls treated with corresponding volume of DMSO in PBS. At E8 eggs treated with EdU as described in the *in ovo* protocol, followed by fixation, immunostaining, and imaging as described above.

### Vocalization analysis

Recordings used for analysis were collected from Cornell Lab of Ornithology’s Macaulay Library. For both call types, three vocalizations were selected per recording and three separate recordings were combined to generate power spectra in Praat, Version 6.4.07 (March 17, 2024)^48^. The male mallard syrinx specimen was provided by the Harvard Museum of Comparative Zoology ornithology department (MCZ:Orn:347156). Scanning with µCT was conducted with X-Tek HMXST225 x-ray imaging system (Nikon Metrology, Inc., Brighton, MI, USA). Acceleration voltage was 60 kV and filament current 180 µA. 3142 projection images captured at 2000 pixels by 2000 pixels. Two frames were averaged with 1 second exposure time per frame. Pixel size = 0.2 mm.

Data reconstructed with Nikon CT Pro 3D software. Bulla volume, neck length, and opening diameter measured using Dragonfly Pro software, Version 2022.2 (Object Research Systems, Inc., Montréal, Canada). Measurements used to model bulla as a Helmholtz resonator:

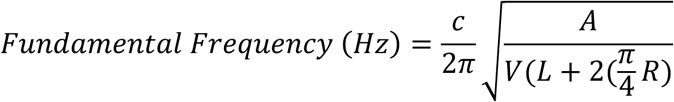

where c = speed of sound (343 m/s), A = neck opening area, V = bulla volume, L = neck length, and R = radius of neck opening area.

This work was performed in part at the Harvard University Center for Nanoscale Systems (CNS); a member of the National Nanotechnology Coordinated Infrastructure Network (NNCI), which is supported by the National Science Foundation under NSF award no. ECCS-2025158.

### Plot generation and statistics (excluding plots from “Image quantification for dorsal and ventral *PITX2* expression”)

Graphing was produced with GraphPad Prism 10 for macOS, Version 10.2.3 (347), April 21, 2024. Normality testing conducted with the Shapiro-Wilk and Kolmogorov-Smirnov tests as well as evaluation of the Normal QQ plot. Parametric statistical testing applied when data showed normal distributions, and non-parametric tests when not. Tests specified in figure legends.

### Single cell RNA-sequencing

#### Sample collection design

10 Experimental groups consisting of combinations of {stage, sex, left/right}. For stage, E6.0, E9.0, E11.0 embryos were prepared, for sex, same representation of male and female, and for each embryo, left and right tissues were collected separately, except for E6.0 stage, where due to the difficulty of splitting the left/right part, the whole tissue was prepared. Each experimental group consisted of exactly one Mallard and one Khaki Campbell breed embryo, and two rounds of sample collection and processing (batch) were performed, such that for each experimental condition, there were four independent embryos. Up to single cell capture, batches were processed on different days, but library preparation, and sequencing were done together.

#### Sample preparation

Tissues were dissected in PBS and placed on ice. Samples incubated on rockers at 37°C with 0.1% trypsin and pipetted until tissue fully dissociated. Cells moved to ice, filtered with 35 µm nylon mesh cell strainer (Falcon 352235), then spun at 200g for 5 minutes at 4°C. Cells were resuspended in PBS with 1% BSA, a small amount reserved for FACS control, then the remainder spun at 200g for 5 minutes at 4°C, and resuspended in DRAQ5 (ThermoScientific, Ref 62251, 5 µM) and DAPI solution (300 nM). The cells were subjected to On-chip Sort microfluidic chip flowshift type sorter. Live cells were collected with DRAQ5+DAPI-gating. Up to this point, the samples were prepared by experimental condition and batch, such that each sample contained one Khaki Campbell and one Mallard embryo. Cell counts for individual experimental groups were determined with a hemocytometer.

#### MULTI-seq labeling & pooling

The 10 experimental groups were labelled with unique MULTI-seq barcodes according to the MULTI-seq protocol^49^. The two batches had all different unique MULTI-seq barcodes. After labelling, still in the cold, all samples were pooled together into a single suspension for each batch and processed through droplet-based single cell capture according to the 10X Chromium Next GEM Single Cell v3.1 (Dual Index) protocol. Subsequent post GEM-RT cleanup, cDNA amplification, and 3’ gene expression library construction steps were performed in accordance with the manufacturer’s instructions, with modification to collect the MULTI-seq barcode, which followed the protocol put forward in McGinnis et al. 2019. Sample indices for the gene expression libraries were chosen from the 10X Dual Index Kit TT Set A (PN-1000215). MULTI-seq library sample indexes were chosen to separate the two batches.

#### Sequencing and preprocessing

Sequencing was performed on a NovaSeq 6000 flowcell at the Biopolymers Facility at Harvard Medical School according to the recommended 10X sequencing specifications. The MULTI-seq barcode libraries were added as well. The sequencing results were obtained in bcl format, demultiplexed by bcl2fastq (2.20.0.422), and the gene expression library fastq files processed via the cellranger pipeline (6.1.0) with the ZJU1.0 genome and gene annotation to obtain the count matrices as well as cellular barcodes. The MULTI-seq libraries were processed with the demultiplex package (https://github.com/chris-mcginnis-ucsf/MULTI-seq).

#### Demultiplexing individual embryonic origin

The cells were demultiplexed in parallel by genotype-based demultiplexing using the vireo package^50^ to ascertain the origin by individual embryos and by the demultiplex package to ascertain the experimental group origin. Based on the genotypic distance between embryos, a clear pattern of two groups was found and with further variant comparison to other in-house datasets from Mallard scRNA-seq, the strain was determined. The inferred doublets from genotypes were discarded in downstream processing. At the same time, the demultiplexed MULTI-seq barcodes were used to associate cellular barcodes by experimental groups. Inferred doublets from MULTI-seq barcodes were also removed for down-stream processing. The associations confirmed that for each MULTI-seq barcode, two vireo-inferred genotypes were found, as expected from the experimental design. It was found that all MULTI-seq barcodes except for the second batch female E9 left experimental condition were sufficiently labelled with similar cellular contributions. Based on the embryonic genotype, except for the left and right condition, The stage and sex was known, so the left/right annotation for the ambiguous cells were labelled and used for downstream processing where left/right difference was not important (e.g. cell type clustering). Lastly, based on the pseudobulk expression profile of W chromosome and Z chromosome by embryo, the individual sex was called (Presence of W chromosome genes and half median expression of Z gene compared to other embryos corresponds to female, absence of W chromosome gene expression and double level of Z gene expression corresponds to males, given the clear absence of sex chromosome dosage compensation in avian species at the RNA level) which was consistent with the MULTI-seq barcode designations, further supporting successful deconvolution of cells into experimental groups at embryo level.

#### Data processing & analysis

The filtered single-cell count matrices were turned into Seurat objects, and all downstream tasks including normalization, cell-cycle phase inference, batch integration, graph-based clustering, and non-linear dimensional reduction visualization were performed using the Seurat v4 framework^51–55^. Leiden algorithm was used for cluster generation^56^, and broad cell type annotation was done manually utilizing well known markers for different cell types. The mesenchymal cluster was used for downstream tasks of differential gene expression analysis. Pseudo-bulk differential gene expression analysis was performed using the deconvoluted cells by embryo using the glmGamPoi package^57^. Design formulas for differential expression accounted for stage and sex.

### 10x single nucleus Multiome ATAC and gene expression

#### Sample collection design

Samples were collected according to the stage and left/right origin. For duck, HH13, HH16, HH17, HH21, and HH25 samples were collected from Mallard and Khaki Campbell breeds. For chick, HH13, HH15/16, and HH25 samples were collected from standard and GFP breeds. Experimental conditions were partially pooled together to carefully balance the left right origin, stage, but such that after demultiplexing the individual embryonic genotype, part of the experimental condition could be inferred back.

#### Sample preparation

Samples prepared following the published protocol with the following exceptions. During nuclei isolation, 10–12 samples pooled for each preparation and pestled for 30 seconds before proceeding to 10-minute lysis. The isolated nuclei were counted with a hemacytometer and then pooled together according to the sample collection design for transposition reaction. Then the nuclei were processed through droplet-based single cell capture according to the 10X Chromium Next GEM Single Cell Multiome ATAC + Gene expression v1 (Dual Index) protocol. Subsequent post GEM-RT cleanup, ATAC library amplification, cDNA amplification and 3’ gene expression library construction steps were performed in accordance with the manufacturer’s instructions. Sample indices for the gene expression libraries were chosen from the 10X Dual Index Kit TT Set A (PN-1000215). Sample indices for the ATAC libraries were chosen from the 10X Dual index Kit NN Set A (PN-1000243).

#### Sequencing and preprocessing

Sequencing was performed on a NovaSeq 6000 flowcell at the Biopolymers Facility at Harvard Medical School according to the recommended 10X sequencing specifications. The sequencing results were obtained in bcl format, demultiplexed by bcl2fastq (2.20.0.422), and the ATAC as well as gene expression library fastq files processed via the cellranger-ARC pipeline (2.0.0) with the ZJU1.0 genome (GCF_015476345.1) and gene annotation for duck (GCF_015476345.1 where the PITX2 gene body was extended 3’ by 700bp to incorporate more reads at the 3’UTR region of the gene), and Galgal6a (NCBI) genome and gene annotation for chick to obtain the count matrices as well as cellular barcodes.

#### Demultiplexing individual embryonic origin

Demultiplexing of individual origin was performed according to the vireo package^50^ with modifications. Briefly, the individual library bam files were modified by cellular barcode suffixes and grouped together for the libraries that partially shared the embryos (left or right origin) for efficient genotyping. For groups that had many cellular barcodes, the cellular barcodes were partitioned into smaller batches and processed, then the resulting inferred genotypes stitched together to ascertain the embryonic origin. The genetic distance was used to ascertain Khaki Campbell or Mallard origin whenever possible. The sex was determined based on the presence or absence of W chromosome genes and Z chromosome gene expression. All cellular barcodes that were inferred to be singlets were subject to graph-based clustering, and mesenchymal cells were selected and pseudo-bulked for Principal Component Analysis (PCA), controlled by sex genotype. The resulting PCA plot shows good correlation of the stages, where the deconvoluted genotypes could be used to infer the stages. Left and right condition could be inferred from the interlocking experimental design for most cases, for those that were ambiguous, the PITX2 expression level in the pseudobulk samples were used to assign its origin.

#### Data processing & analysis

The filtered single-cell count matrices were turned into Seurat objects, and all downstream tasks including normalization, cell-cycle phase inference, batch integration, graph-based clustering, non-linear dimensional reduction visualization was performed using the Seurat v4 framework^51–55^ as well as the Signac package^58^. Leiden algorithm was used for cluster generation^56^, and broad cell type annotation was done manually utilizing well known markers for different cell types. The mesenchymal cluster was used for downstream tasks of differential gene expression analysis. Pseudo-bulk differential gene expression analysis was performed using the deconvoluted cells by embryo using the glmGamPoi package^57^. Design formulas for differential expression accounted for stage and sex.

### Duck vocalization and air pressure measurements

Ducks were fitted with a harness around the thorax and wings with a Velcro tab attached on the back and returned to the group of 3 females and 1 additional male. After a few days of getting adjusted, surgical procedures were initiated for measurement of air pressure in the bulla. Ducks were deprived of food and water for 3 hours. Prior to surgery, the skin overlying the pectoralis and clavicular area was treated with procaine gel. After a few minutes, a Ketamine/Xylazine mixture was administered. Once surgical depth was reached, the syrinx was accessed by an incision in the skin, removal of the fat to one side, and opening of the interclavicular air sac membrane. Once the syrinx was exposed, we drilled a small hole into the bulla surface and inserted a stainless steel cannula to which a flexible tube (Dow Corning Silastic tubing) was attached. The cannula was secured to the bulla surface with tissue adhesive (Vetbond) and the tube was guided out of the air sac and routed subcutaneously to the back. The air sac membrane was closed with suture and tissue adhesive, the fat returned to its original position and the skin sutured closed. The free end of the cannula on the back was attached to a pressure transducer (Fujikura FPM-02PG).

After birds were awake and started feeding, they were returned to the group and were allowed 1-2 days of recovery. Then the pressure transducer was plugged in to an amplifier and power supply (Hector Enfgineering, custom built box) via a long cable. To avoid birds getting tangled in the wires, a tether system lifted these up and allowed free movement of the instrumented individual. The pressure signal was recorded together with sound (AudioTechnica AT8356) at 44.1 kHz using Avisoft Recorder software. Recordings were analyzed with Praat software. All procedures were approved by the IACUC of the University of Utah.

## Acknowledgements

We thank Jeremiah Trimble and Katherine Eldridge from the Harvard Museum of Comparative Zoology Ornithology Collections. Hao-Yu Greg Lin at the Center for Nanoscale systems for help with µCT. Paula Montero Llopis, Praju Vikas Anekal, and Adrienne Wells for training and aid with microscopy. Chad Araneo for duck eggs.

## Notes

### Competing Interest Statement

The authors have declared no competing interest.

