## Supplemental Figures for "Integration of sexual dimorphism and left-right asymmetry in the development of the duck syrinx"

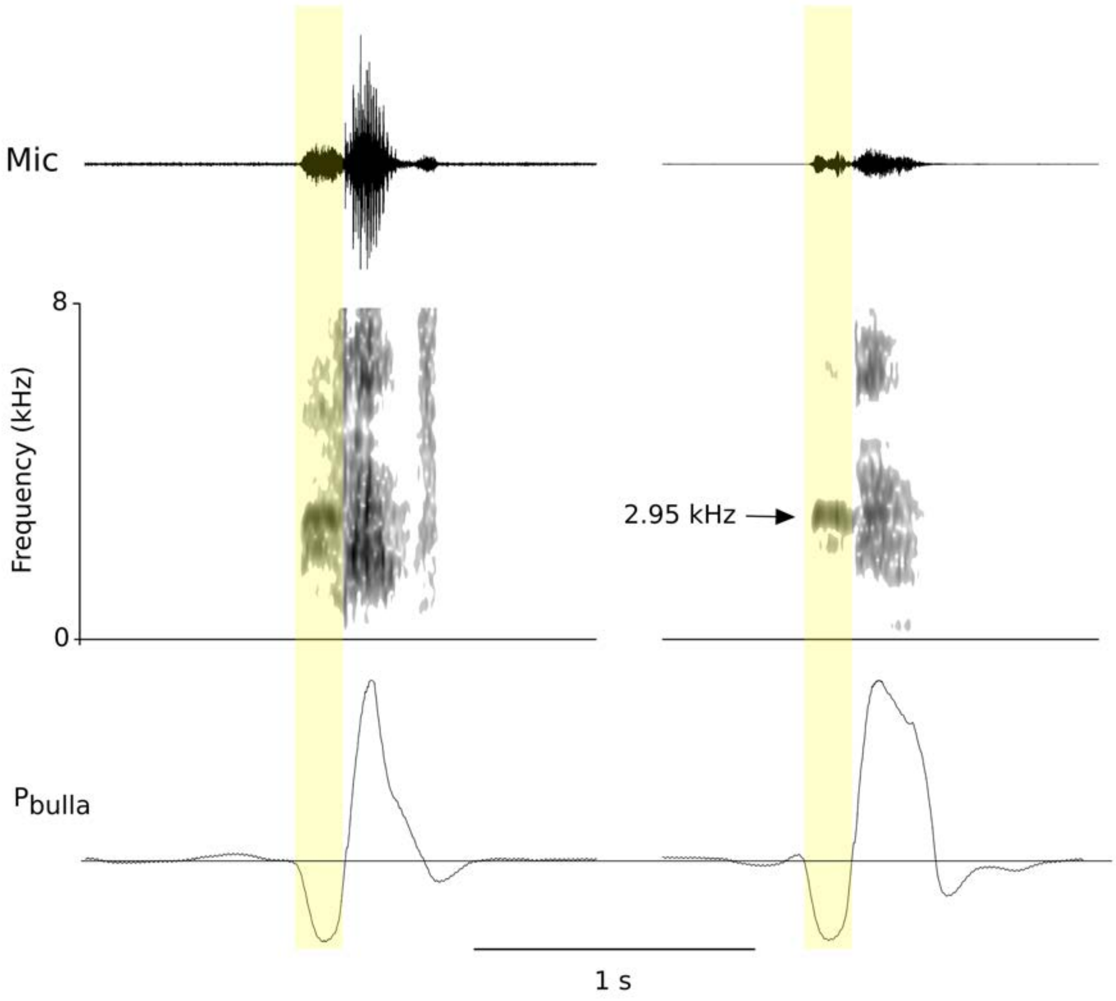


**Figure S1. High frequency vocalization in Mallard occurs on the inspiration.**
(A) Ventral view of µCT scan of Mallard syrinx. Scale bar 1cm. (B) Recording of Mallard vocalizing. Top, sound track; middle, sonogram; bottom, pressure in the bulla. Yellow highlight shows region with fundamental frequency matching that of the courtship whistle. Pressure drops as this occurs, indicating the vocalization is on the inspiration. Scale bar 1 second.

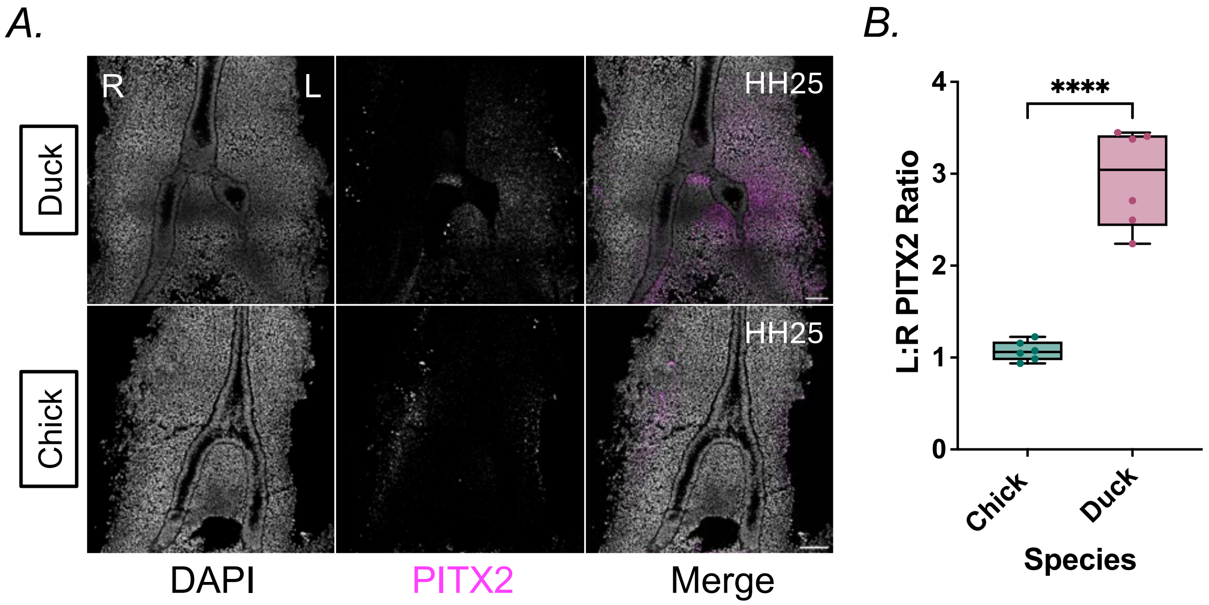


**Figure S2. Comparison of *PITX2* expression shows left-right asymmetry in duck, not chick.** (A) PITX2 HCR in duck (above) and chick (below) at HH25. Probes generated against all PITX2 isoforms; PITX2a and PITX2b expected bilaterally while PITX2c is expected to be asymmetric. (B) Quantification of Pitx2 expression. Individual RNA transcripts were counted using the Matlab plugin ImageM. The ratio of RNA transcripts/total cell number (DAPI) on the left vs. the right was collected for each biological replicate. Chick n = 6, duck n = 6. Two-tailed Student's t-test, **** $p < 0.0001$.

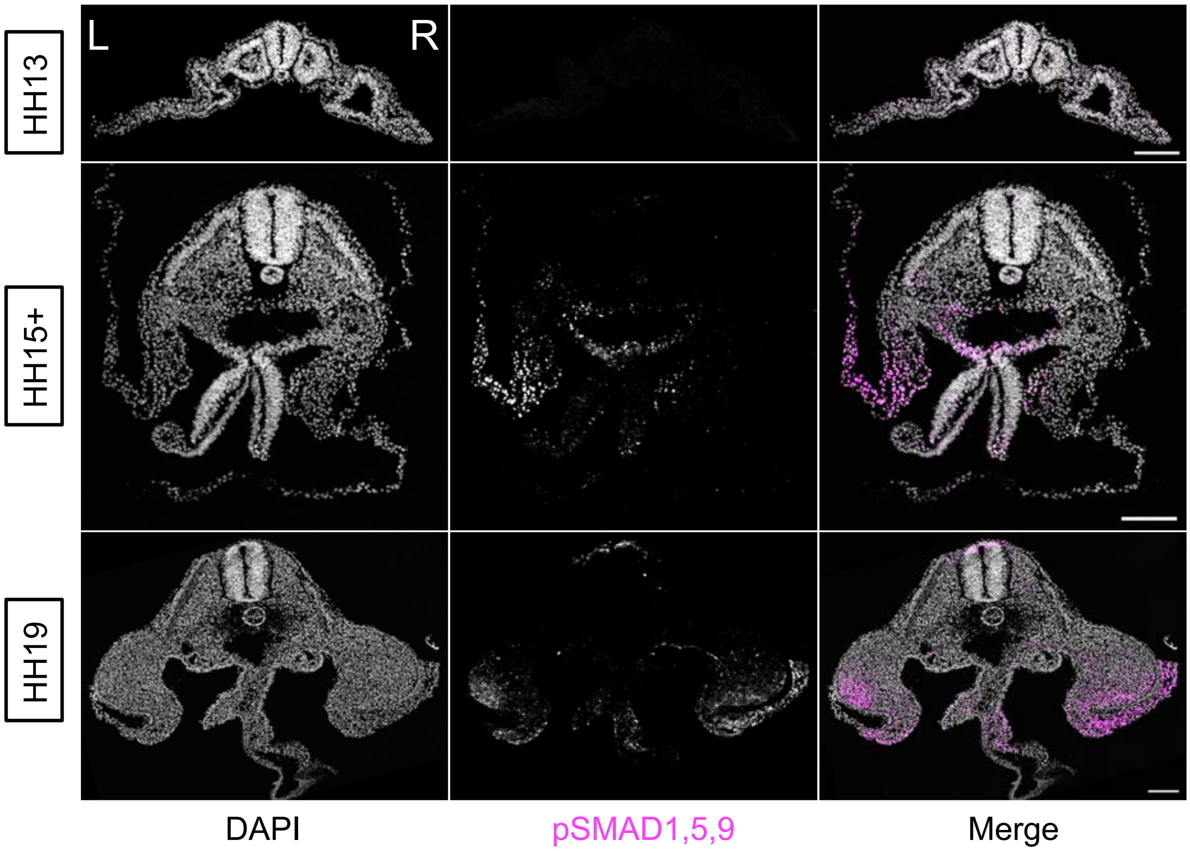


**Figure S3. BMP signaling is active in the chick embryo.**
Maximum intensity projections of representative HH13 (3/3), HH15+ (4/4), and HH19 (3/3) chick embryos stained with pSMAD1,5,9. Right (R) and left (L) as labeled on image. Scale bar = 100μm.

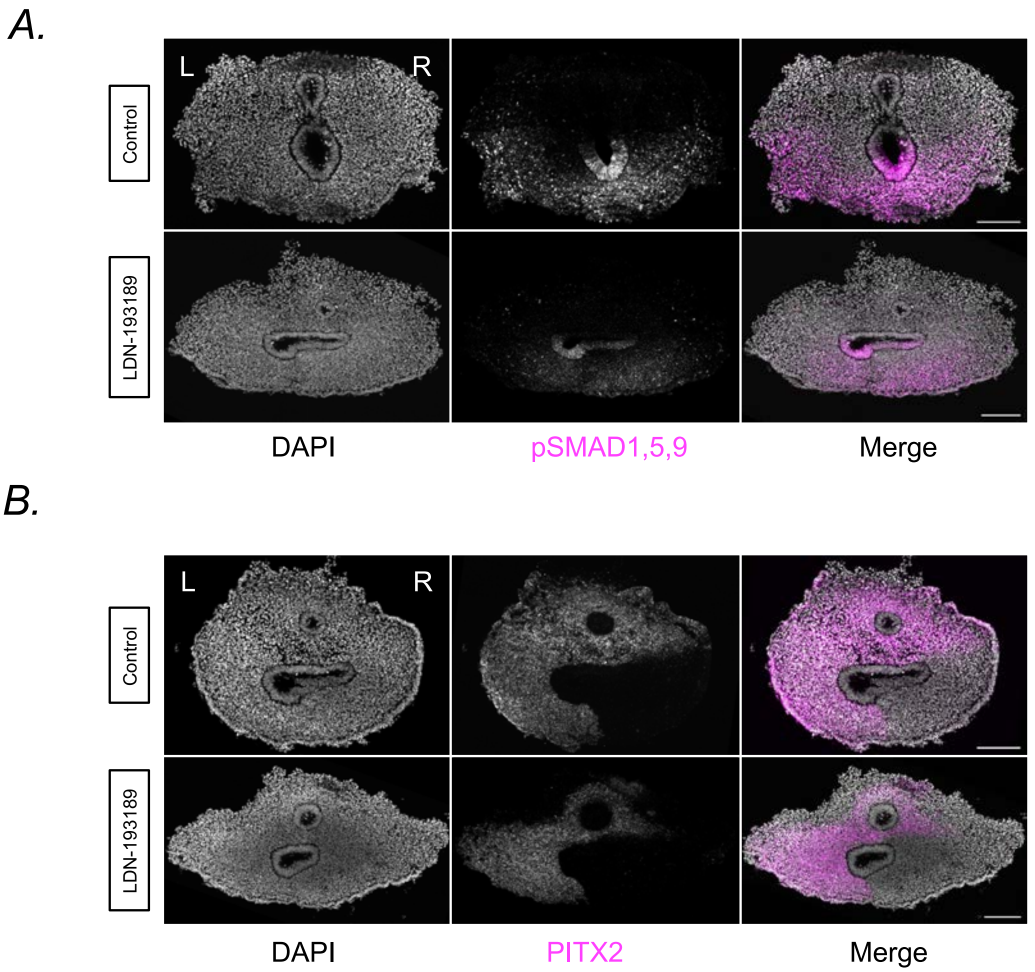


**Figure S4. Controls for LDN-193189 treatment showing BMP inhibition with LDN-193189 results in reduced *PITX2* expression.**
(A) Maximum intensity projection of control (above) vs. LDN-193189-treated (below) foregut at HH21 shows reduction of pSMAD1,5,9 immunofluorescence after chemical treatment with LDN-193189 (0.375mM per egg). (B) Maximum intensity projection of HCR for PITX2 in control (above) vs. LDN-193189-treated (below) embryos. Right (R) and left (L) as labeled on image. Scale bar = 100µm.

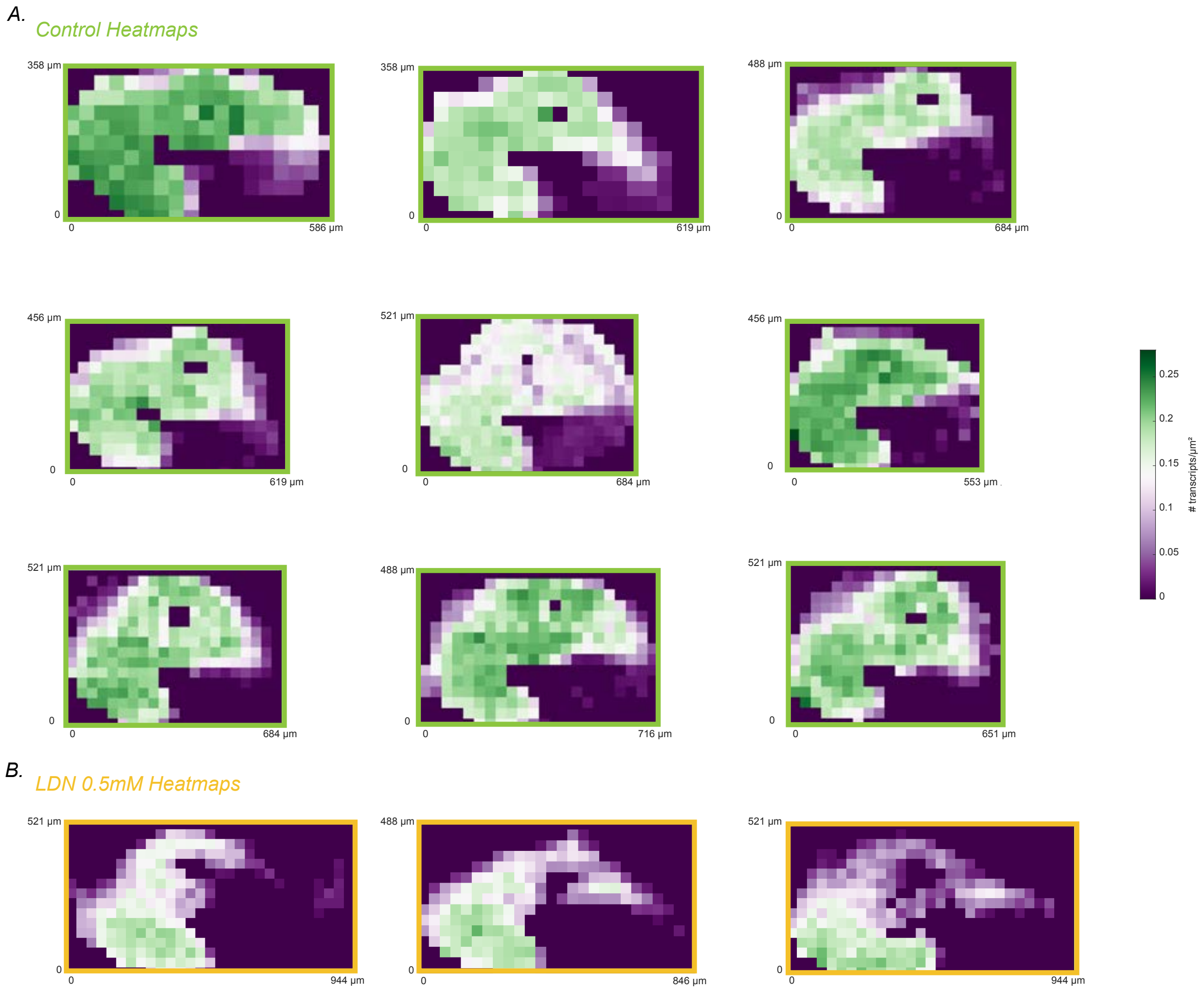


**Figure S5. Heatmaps of *PITX2* expression for quantified samples not visualized in the main text.** (A) Heatmaps of control samples outlined in green. (B) Heatmaps of samples treated with 0.5mM LDN-193189 outlined in yellow.

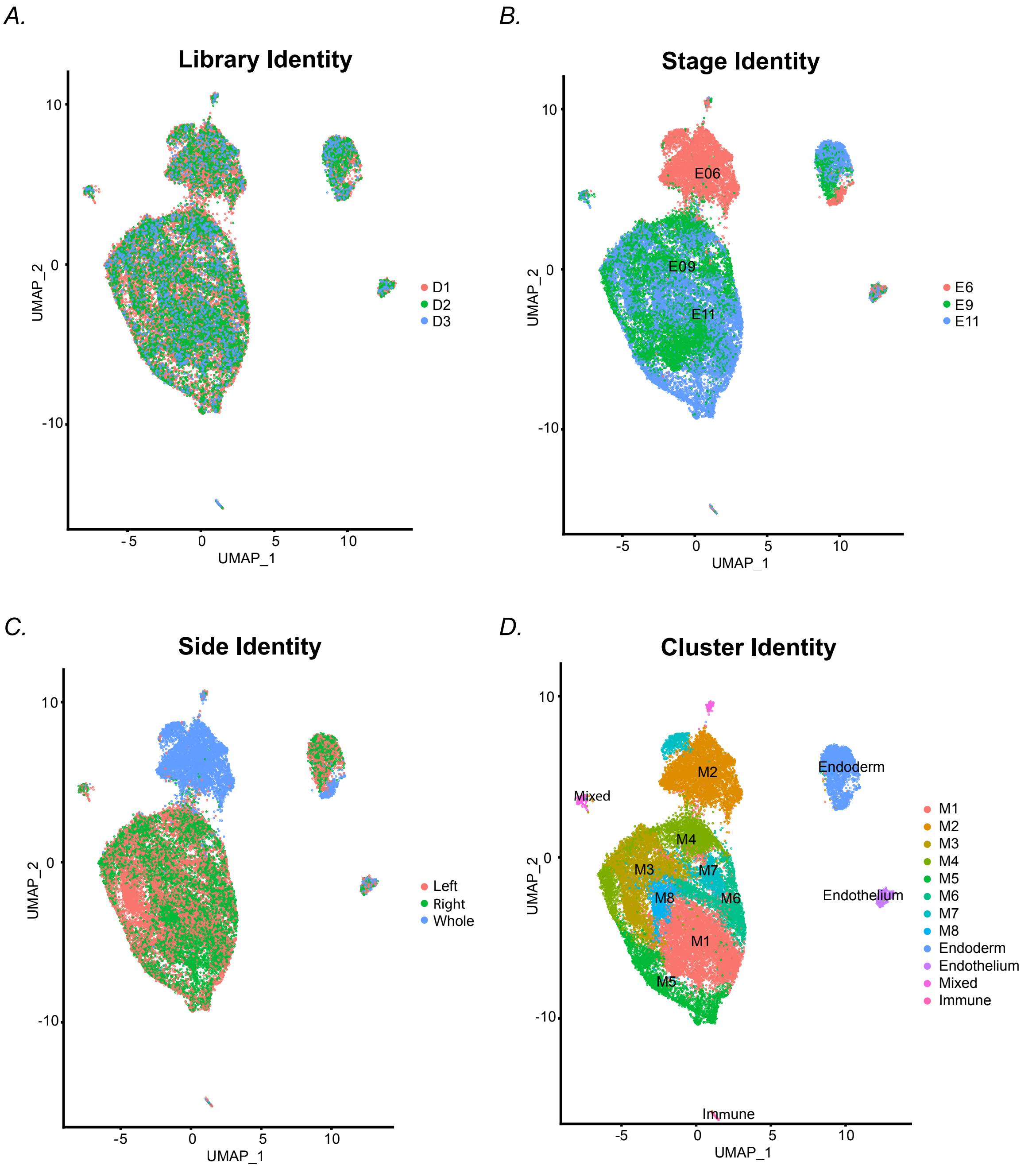


**Figure S6. UMAP plots of scRNA-seq dataset**. Distribution shown based on (A) library, (B) developmental stage, (C) tissue side (note E6 tissue not divided into halves), or (D) cluster identity. Clusters M1–M8 are mesoderm clusters 1–8. Mixed refers to a population including fibroblasts, smooth muscle, and immune cells.

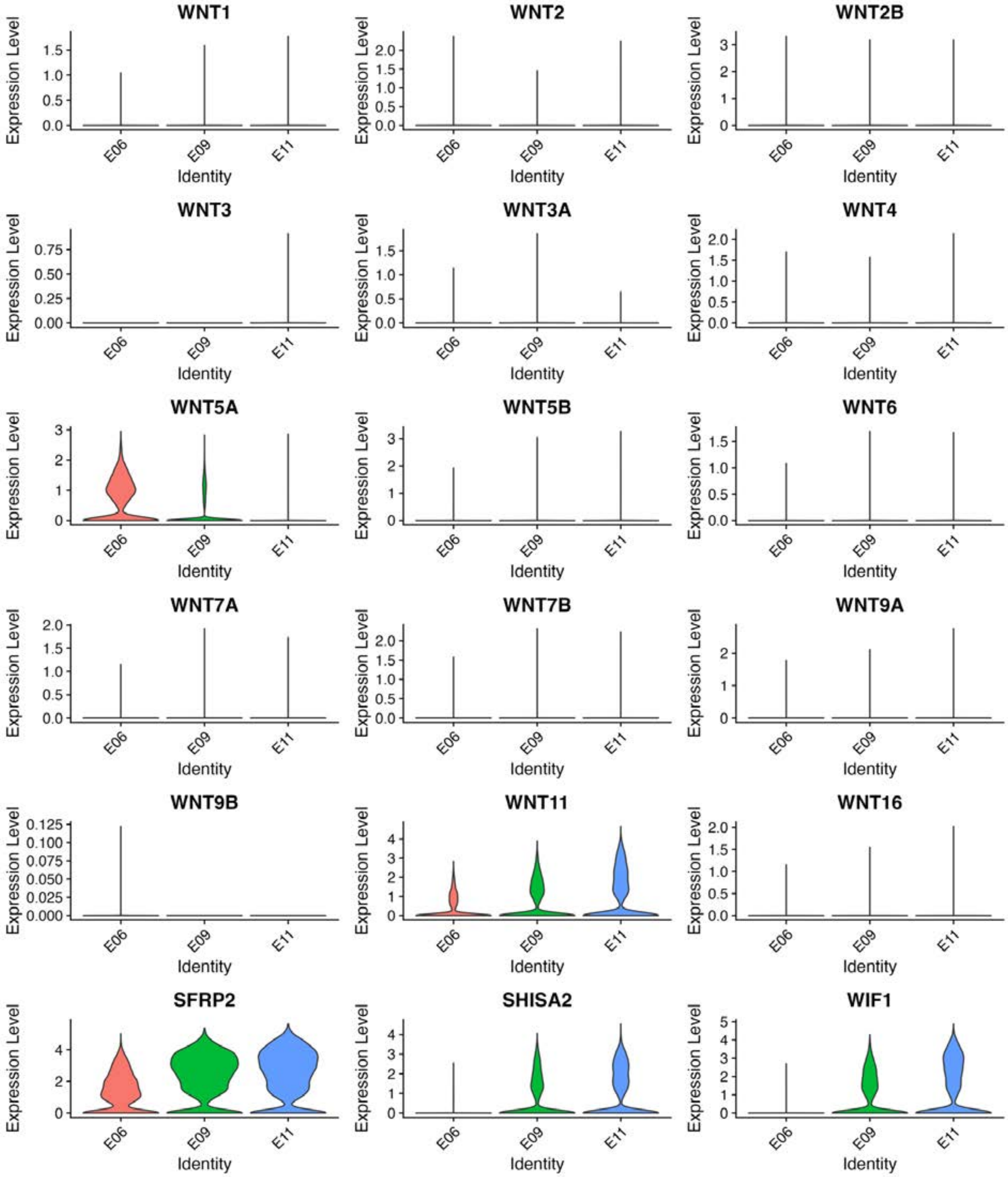


**Figure S7. Violin plots showing expression of WNT ligands and pathway modulators.** Plots show RNA expression in pooled mesoderm clusters over developmental time.

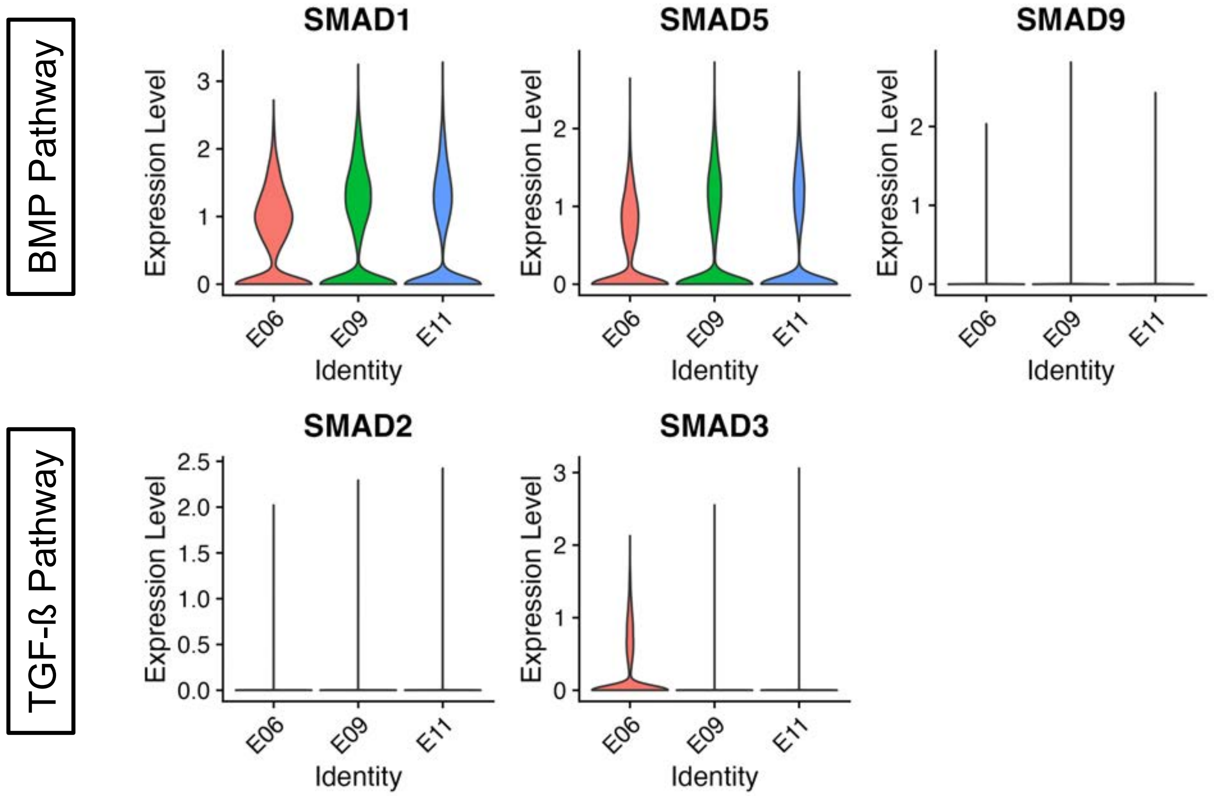


**Figure S8. The BMP pathway, not the TGF-ß pathway is active in the syrinx**. Violin plots of genes in the BMP pathway (above) and TGF-ß pathway (below) for pooled mesoderm clusters over time.

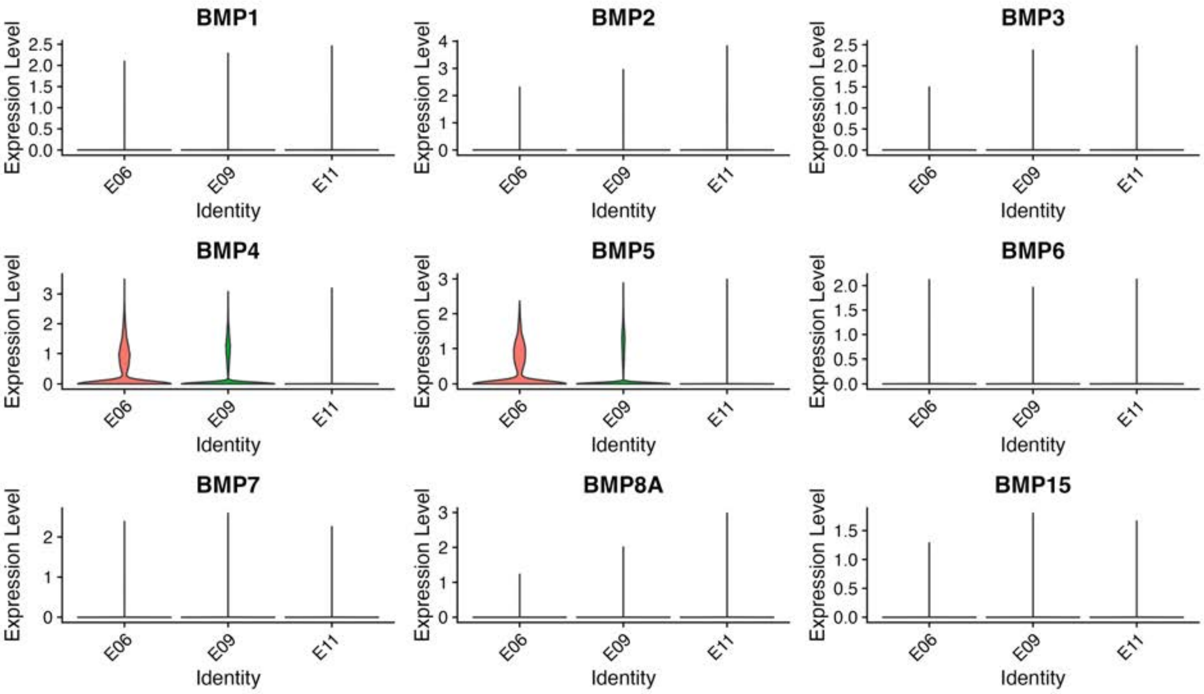


**Figure S9. Violin plots showing expression of BMP ligands and pathway modulators.** Plots show RNA expression in pooled mesoderm clusters over developmental time.

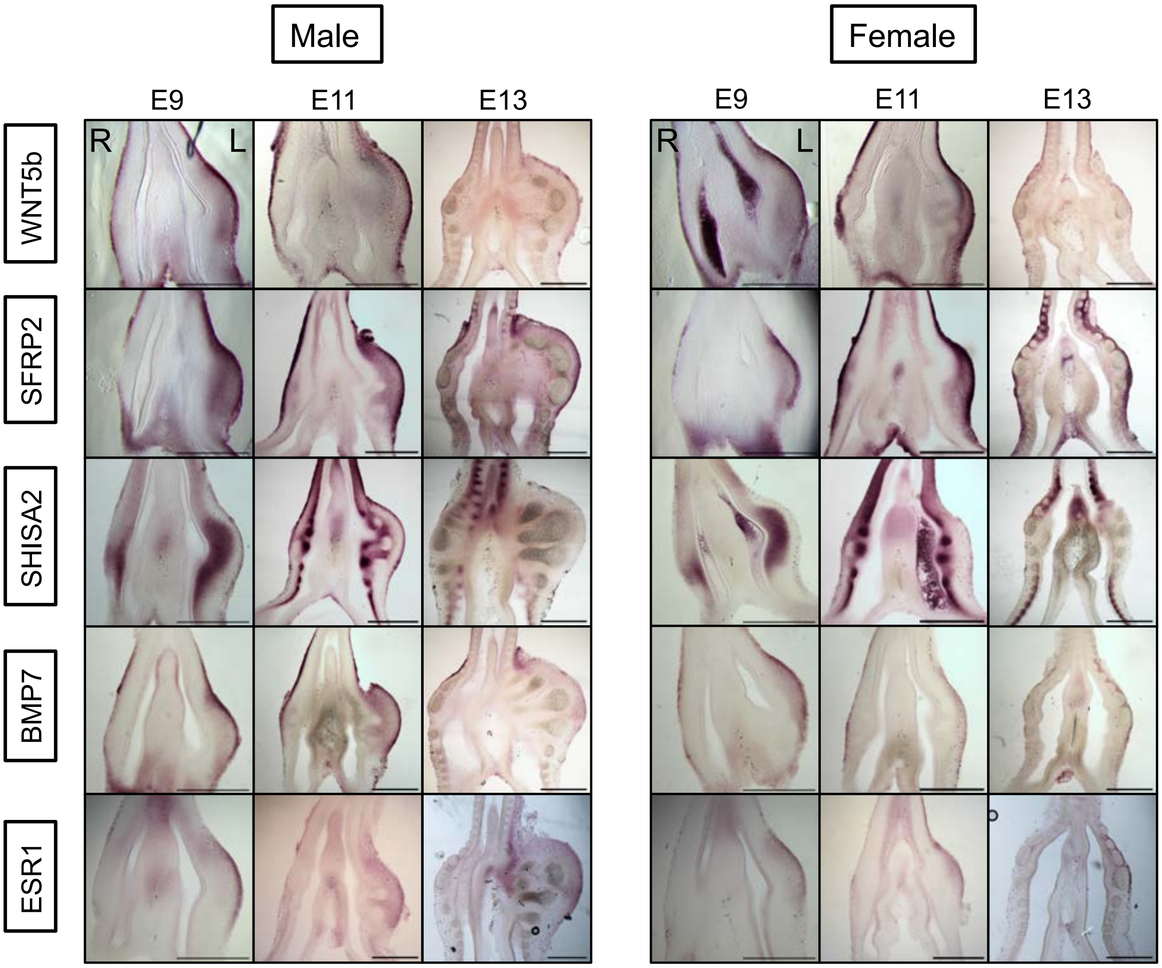


**Figure S10. *In situ* hybridizations of gene candidates generated from the scRNA-seq dataset.** Left, male E9, E11, and E13 *in situ* hybridizations of WNT5b, SFRP2, SHISA2, BMP7, and ESR1. E9 images reproduced here from Figures 5b and 6b of the main text for comparison. Right, female E9, E11, and E13 *in situ* hybridizations of WNT5b, SFRP2, SHISA2, BMP7, and ESR1. Right (R) and left (L) as labeled on image. Scale bar = 0.5mm.
